# Global patterns and environmental drivers of soil fungal nematode antagonists

**DOI:** 10.1101/2025.11.12.688024

**Authors:** Robbert A.F. van Himbeeck, Geert Smant, Julian Helftenstein, Stefan Geisen, Johannes Helder

## Abstract

Plant-parasitic nematodes can be suppressed by antagonistic members of the local soil microbiome. Compared to bacterial antagonists, considerably more attention has been paid to nematode-suppressing fungi. However, little is known about their global distribution. As their biogeography matters from both a fundamental and an applied perspective, we set out to mine the GlobalFungi database. After filtering on sample type, biome relevance and sequencing depth, we retained approximately 28,000 samples from 484 studies. An overview of fungal nematode antagonists was generated, and subsequent analyses revealed that 82.6% of the soil samples comprised ≥1 nematode antagonist. Most of these antagonists are not obligatory nematode parasites; they switch between trophic lifestyles including saprophytism. Nematode antagonists are frequently found across most biomes; the detection probability was highest in croplands (88% of samples), woodlands (82%), and grasslands (76%). Most of the common nematode antagonists are present on all continents and across multiple climate zones underlining their enormous ecological flexibility. With one exception, the most frequently detected antagonistic fungi belonged to the fungal order Hypocreales whose members are known to parasitize insects and fungi as well. Analysis of the impact of temperature and precipitation on common antagonists in croplands revealed mostly non-linear responses. Among a selection of six soil properties, pH was the most informative predictor for the abundance of antagonists in croplands. Insights into the prevalence and the distribution of nematode antagonists at a global scale contribute to the exploration of the nematode-suppressive potential, which appears to be more common and widespread than often assumed.

## 1. Introduction

Soil-borne plant diseases constitute a major threat to global crop production, and their global prevalence is projected to increase due to global warming (Delgado-Baquerizo et al., 2020). Fungi, bacteria, viruses, oomycetes, and nematodes cause soil-borne diseases, and the most harmful pathogens are typically controlled by measures such as crop rotation, the cultivation of resistant varieties, and the application of agrochemicals. However, many pathogens are polyphagous, for numerous crops no resistant varieties are available, and for environmental reasons the use of agrochemicals is more-and-more restricted. Natural disease-suppressiveness is another category of plant protection where plant diseases are suppressed by the local soil microbiome (Schlatter et al., 2017). This phenomenon has been described for various soil-borne plant diseases, such as *Ralstonia solanacearum* (Su et al., 2022) *Pythium myriotylum* (Oni et al. 2020), and *Fusarium culmorum* (Ossowicki et al., 2020). Plant-parasitic nematodes, invertebrate pests that affect all major staple crops, have also been observed to be suppressed by specific microbial antagonists (Topalovíc et al., 2020). Suppressive soils have been identified for species such as *Meloidogyne javanica* (Watson et al., 2020), *Globodera pallida* (van Himbeeck et al., 2025), *Heterodera avenae* (Kerry et al., 1982), and soybean cyst nematodes (Hu et al., 2017) along with their associated bacterial and fungal antagonists. Kerry et al. (1982) observed a natural decline in *H. avenae* under continuous cropping of susceptible cereals, which they could attribute to native populations of *Nematophthora gynophila* (Oomycota) and *Metacordyceps* (anamorph: *Pochonia*) *chlamydosporia* (Fungi). Although both bacterial and fungal members of soil communities can suppress nematodes, fungi have historically received most attention, largely due to their ability to produce pronounced and diverse nematode-trapping devices (*e.g.,* Pramer, 1964).

Some fungi antagonize nematodes directly by trapping mobile life stages of the nematode (*e.g., Arthrobotrys* spp.; Scholler et al., 1999), by parasitizing eggs or females (*e.g., Metacordyceps chlamydosporia*; Manzanilla-López et al., 2013), by the release of toxins (*e.g., Pleuroteus ostreatus*; Nyangwire et al., 2024), or by endoparasitism (*e.g., Hirsutella rhossiliensis*; Hallmann et al., 2019). Many direct antagonists are facultative parasites and their nematophagous activity primarily relies on the nutrient status of the soil (Baweja and Rawat, 2020; Luambano et al., 2015), the presence of nematodes (Baweja and Rawat, 2020; Nordbring-Hertz, 1968), and edaphic conditions such as pH and soil texture (Mo et al., 2005). The nutrient-rich conditions in most arable soils likely promotes a saprophytic rather than a nematophagous lifestyle among many facultative nematophagous fungi. Another category of fungi antagonizes nematodes indirectly by eliciting induced systemic resistance (ISR) (Zheng et al., 2024) and/or systemic acquired resistance (SAR) (Salwan et al., 2022) in plants, thereby priming the plants’ defence system against pathogen infections. Fungal biocontrol agents have been developed against plant-parasitic nematodes (Li et al., 2015), often showing substantial suppression under highly controlled conditions (Bairwa et al., 2023; Tazi et al., 2021; Zhang et al., 2020). However, the success of these products under field conditions remains limited and inconsistent (Ahmad et al., 2021; Albright et al., 2022).

Nematode antagonists occur in a wide variety of soil types worldwide ranging from forest soils (*e.g.,* Rong et al., 2020) to intensively managed croplands (*e.g.,* van Himbeeck et al., 2025). Certain antagonists, such as *Arthrobotrys oligospora*, have been detected frequently in soils (*e.g.,* Deng et al., 2023b; Doolotkeldieva et al., 2021; Berhanu et al., 2022), while others are rarely encountered. This is illustrated by a recent survey of nematode-trapping fungi across Yunnan Province, China, where *A. oligospora* was detected in 190 of the 228 sampling sites, whereas various *Dactylellina* and *Drechslerella* species were observed only at a single site (Deng et al., 2023a). The importance of abiotic soil properties such as pH (Tedersoo et al., 2020; Glassman et al., 2017) in shaping fungal communities is well-known, but edaphic properties driving nematode-antagonistic fungal diversity are poorly understood. Stimulation of the abundance and activity of fungal nematode antagonists by organic amendments has been reported frequently (Jaffee, 2004; Stirling et al., 2011; Rosskopf et al., 2020), but the influence of physicochemical soil properties remains largely unknown. Most soil fungi have a broad global distribution that is predominantly governed by environmental conditions such as mean annual temperature and precipitation (Tedersoo et al., 2014; Větrovský et al., 2019). We anticipate that edaphic and climatic conditions also drive global distributions of fungal nematode antagonists. However, despite of their frequent detection, the biogeography of fungal nematode antagonists, including how this is shaped by environmental conditions, has never been documented systematically.

The aim of this paper was to determine the biogeography of fungal nematode antagonists at the global level and to assess how this is affected by environmental conditions. After filtering the GlobalFungi database for relevant biomes and read coverage, we analysed 27,932 samples and focused on known nematode antagonists. Additionally, we linked mean annual temperature (MAT) and annual precipitation (AP) to antagonist presence and relative abundance. For the cropland samples, we further examined how soil properties (*e.g.,* pH and soil organic carbon) were associated with model-predicted antagonist relative abundances. With this approach, we addressed the following research questions: i) How prevalent are fungal nematode antagonists in global soil samples, and which antagonistic taxa are most widespread?; ii) Which biomes and climatic conditions favour the presence and relative abundance of antagonists?; and iii) Which soil properties are most important for predicting antagonist relative abundance in croplands?

## 2. Materials and methods

### 2.1. GlobalFungi dataset, filtering, and antagonist selection

We used the GlobalFungi database (release 5, Větrovský et al., 2020) to assess the global distribution of fungal nematode antagonists. This database comprises Internal Transcribed Spacer (ITS) 1 and ITS2 sequences from fungal communities based on 846 individual studies comprising a total of 84,972 samples collected across the globe. In this database, samples originate from various types (*e.g.,* soil or water) and biomes (*e.g.,* cropland or mangrove). The ITS sequences in the database are centrally and uniformly processed (for details, see Větrovský et al., 2020). For this study, we downloaded the GlobalFungi database containing only species-level annotations, including annotations for both ITS1 and ITS2 sequences. Subsequently, we filtered the entries based on sample type (including “litter”, “rhizosphere soil”, “root”, “root + rhizosphere soil”, “soil” and “topsoil”) and biome (including “anthropogenic”, “cropland”, “desert”, “forest”, “grassland”, “shrubland”, “tundra”, “wetland”, and “woodland”), while manipulated samples were excluded. The biome “anthropogenic” includes areas that are directly disturbed by human activity, such as mining sites, and it is noted that croplands and managed forests are not included in this category. To reduce the chance of false-negative antagonist detections, only samples with >20,000 reads were used for further analyses. After these filtering steps, 27,932 samples from 484 studies were retained (Figure 1A). Of all ITS sequences in the resulting dataset, on average 49.3% was identified till species-level.

**Figure 1:**
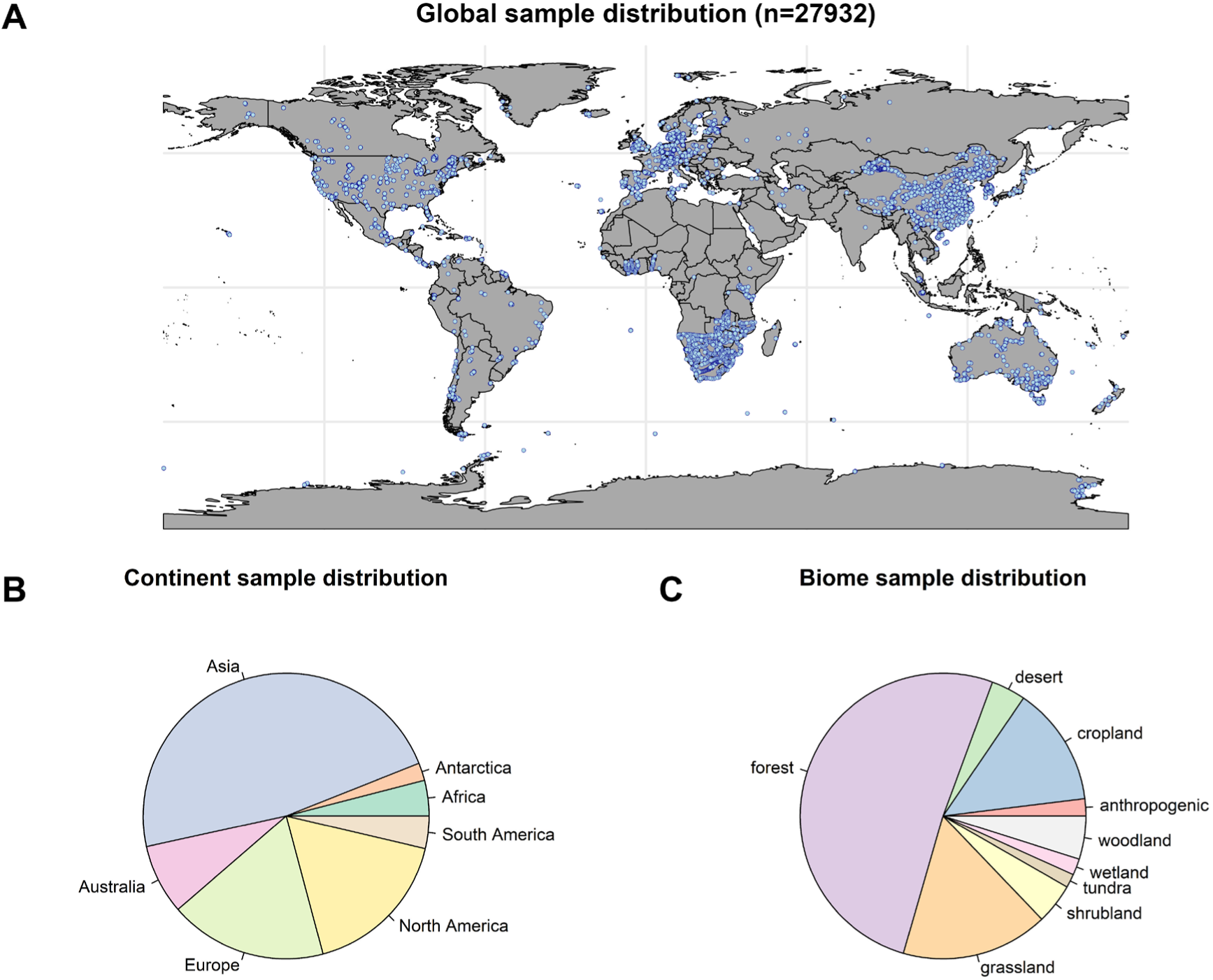
Overview of the sample distribution of the filtered GlobalFungi dataset, consisting of a total of 27,932 samples. A) World map showing the location of all individual samples (blue dots). B) Sample distribution per continent. C) Sample distribution per biome.

To the best of our knowledge, there is no public database of nematode-antagonistic fungi. Therefore, we generated a list of (putative) nematode-antagonists, and in this selection, we included activators of the plant innate immune system that were shown to reduce plant-parasitic nematode establishment. All queried nematode antagonists are provided as R vector in Supplementary Data S1, whereas those that were detected in the GlobalFungi database are listed in Supplementary Table S1. Some strains of the *Fusarium oxysporum* species complex have been described as nematode antagonists (Waweru et al., 2014; Dessel et al., 2011). However, because of the ecological and functional diversity of this taxon (Edel-Hermann and Lecomte, 2019), it was excluded from the current analyses

### 2.2. Data processing and statistical analyses

All analyses were performed in R (v4.3.1) and all plots were generated using ggplot2 (v3.4.2; Wickham, 2016), except for the sample distribution pie charts (base R). Singletons were excluded from all analyses. As the presence and abundance of native nematode antagonists is of particular interest in croplands, we placed special emphasis on this biome.

#### 2.2.1 Visualization of sample occurrence and global distribution

Barplots were created to visualize the occurrence (in percentage of samples) of detected antagonists. Geospatial data processing and visualization were performed using the sf (1.0.19; Pebesma and Bivand, 2023) and rnaturalearth (v1.0.1; Massicotte and South, 2023) packages, in combination with ggplot2. For these plots, the six most common nematode-antagonistic taxa in croplands, namely *Clonostachys rosea*, *Acremonium persicinum*, *Chaetomium globosum*, *Metarhizium anisopliae*, *Trichoderma asperellum*, and *Beauveria bassiana*, were selected. Additionally, the distribution of a number of nematode-antagonistic fungi that are applied as biological control agents in croplands was visualized: *Metacordyceps chlamydosporia*, *Purpureocillium lilacinum*, *Trichoderma harzianum*, *Trichoderma viride*, *Arthrobotrys* spp., and *Hirsutella rhossiliensis*. The fungi *M. chlamydosporia, P. lilacinum, T. harzianum, T. viride* and *Arthrobotrys* spp. are included in various formulations used to control root-knot nematodes (Li et al., 2015; Groves et al., 2025; Mukhtar et al., 2021), and *H. rhossiliensis* is applied to control beet cyst nematodes (Hallmann et al., 2019). In this study, the terms ‘common’, ‘prevalent’, ‘frequently observed’, and ‘rare’ refer to the frequency with which a taxon was detected across samples, irrespective of its distribution across the globe. On the other hand, the terms ‘widespread’, ‘distribution,’ and ‘biogeography’ refer to the spread of taxa across the globe.

#### 2.2.2 Mode of antagonism

Based on literature (see Supplementary Table S1 and Topalovíc et al., 2020; Li et al., 2015) the detected antagonists were divided according to their principal mode of antagonism: Egg & female parasitic, nematode-trapping, endoparasitic, toxin-producing, and multiple, including induced systemic resistance (ISR) (Supplementary Table S1). For some taxa, such as the antagonistic members of the genus *Trichoderma*, multiple modes are described and this regularly includes non-nematode specific antagonism by general stimulation of the plant immune system (*i.e.,* ISR). Therefore, these taxa were classified as ‘Multiple, incl. ISR’.

#### 2.2.3 Patterns of antagonist presence and relative abundance across biomes and climatic conditions

##### Climatic data acquisition and data preprocessing

Global mean annual temperature (MAT) and annual precipitation (AP) with a 30 arcseconds resolution were retrieved from the WorldClim database (version 2.1) using the geodata (v0.6.2; Hijmans et al., 2024) package. Next, the terra package (v1.8.29; Hijmans, 2025) was used to rasterize the climatic variable data and to extract the climatic data for all GlobalFungi sample coordinates. Prior to statistical modelling, we curated the dataset to ensure consistency of the predictor variables. Samples for which no climatic data could be extracted were excluded from the dataset, as these variables were central to our hypothesis and included as fixed effects in the models. Additionally, we removed samples for which the ‘month of sample’ variable spanned more than one month (*e.g.,* January-April), to ensure temporal consistency across samples. These filtering steps resulted in a total of 19,450 samples remaining for the subsequent statistical analyses). For all models, the climatic variables were scaled (μ = 0, SD = 1) to improve model fit. As the nematode trapping capacity of *Arthrobotrys* is conserved across the genus, aggregated data of *Arthrobotrys* species were analyzed.

##### Modelling antagonist presence

The effect of biome and climatic variables (*i.e.,* MAT and AP) on antagonist presence was modelled using a Generalized Linear Mixed Model (GLMM) with binomial distribution (package: lme4 v1.1.34; Bates et al., 2015). Antagonist presence was included as response variable, while biome, MAT, and AP were included as fixed effects. To account for potential non-linear responses to climatic variables, quadratic terms of MAT and AP were also included. Variables ‘month of sampling’, ‘continent’, and ‘paper_ID’ were included as random effects. For each biome, population-level predicted probabilities of antagonist presence were obtained from the model and averaged across observations. Within-biome variation in predicted probabilities across climatic conditions was visualized using the 2.5^th^ and 97.5^th^ percentiles of population-level model predictions and displayed as error bars.

##### Modelling antagonist relative abundance

The effect of biome and climatic variables on antagonist relative abundances was modelled using GLMMs with negative binomial distribution. The response variables were defined as raw taxon read counts of (i) the aggregated antagonist data (*i.e.,* all antagonist reads summed), (ii) the six most common cropland taxa, and (iii) the six taxa frequently applied as biological control agents. To account for variation in sequencing depth, the logarithm of the sample sequencing depth was added as offset in all GLMMs addressing antagonist relative abundance. This approach allows modelling of relative abundances while retaining count-based statistical properties (Zhang et al., 2017; Cazzaniga et al., 2025). Biome, MAT, and AP were included as fixed effects and quadratic terms of MAT and AP were also included to account for potential non-linear responses to climatic variables. Also for this analysis, variables ‘month of sampling’, ‘continent’, and ‘paper_ID’ were included as random effects. For the models of *Trichoderma asperellum*, *Metacordyceps chlamydosporia* and *Hirsutella rhossiliensis*, ‘continent’ was excluded as random effect because its estimated variance was approximately zero.

##### Model diagnostics

DHARMa (v0.4.6; Hartig, 2022) Q-Q and residuals vs. predicted values plots revealed proper model fit for all GLMMs, except for the negative binomial model fitted to the aggregated antagonist dataset. For this model, deviations from residual uniformity were observed, which likely reflect heterogeneity in responses among antagonist taxa that could not be captured by a single global model. However, no evidence of dispersion issues was observed and hence the model was retained. Zero-inflation of the models was assessed using the performance package (v0.13.0; Lüdecke et al., 2021). A slight overprediction of zeros was detected in two models (the aggregated antagonist dataset and *Clonostachys rosea*), but this deviation was minor and did not affect model interpretation.

#### 2.2.4 Impact of various soil properties on antagonist relative abundance in croplands

##### Soil data acquisition and preprocessing

Various soil properties can be modified in croplands through farm management practices (*e.g.,* pH by liming), making them potential targets for manipulating native nematode-antagonistic fungal communities. Thus, following the global analysis of biome and climatic effects, we used random forest models on the cropland subset (n = 3,162) of this dataset to explore the relative importance of local soil properties in explaining antagonist relative abundances in croplands across continents. For this, predicted mean values of soil pH, nitrogen, cation exchange capacity (CEC), bulk density (bdod), coarse fragment content (cfvo; volumetric percentage of particles >2 mm), organic carbon density (ocd), soil organic carbon (soc), and soil texture (sand, silt, and clay as mass percentage) were extracted for each cropland sample from the SoilGrids2.0 database (Poggio et al., 2021) using the sf (v1.0.23; Pebesma, 2018) and terra (v1.8.86; Hijmans, 2020) packages. For each sample we calculated the overlap of its sampling depth with the six SoilGrids depth layers (Poggio et al., 2021) and calculated a thickness-weighted average of the overlapping layers. Subsequently, the resulting weighted averages were divided by their respective conversion factor (Poggio et al., 2021) to recover the true values. For 393 (12.5%) out of the 3162 selected cropland samples, no sampling depth was provided. For these samples, a sampling depth of 0-30 cm was assumed, as sampling between 0-30 cm is very common for soil sample collection in croplands (*e.g.,* Hannula et al., 2021; van Himbeeck et al., 2025) and 92% of the cropland samples in this study originated from this soil layer. Samples without overlap with Soil Grids layers (n=44; 1.4%) or no retrievable SoilGrids data (n = 283; 8.9%) were excluded. This resulted in a total of 2,835 (89.6 %) cropland samples being included in the subsequent analyses. To account for multicollinearity among soil property variables, one variable of variable pairs showing an absolute correlation greater than 0.7 was excluded from the analysis. As a result, the variables nitrogen, bdod, and silt were excluded.

##### Statistical modelling and interpretation

We fit random forests regression models using the randomForest package (v4.7.1.2;Liaw & Wiener, 2002) (ntree=500, mtry=p/3, where p is the number of predictors) to model the associations between the soil property data (as predictor variables) and the relative abundance of (i) the aggregated antagonist data, (ii) the six most common cropland taxa, and (iii) the six taxa frequently applied as biological control agents. Only random forest models that explained >10% of the variation were used for further analyses. After fitting random forest models, we used the treeshap package (v0.3.1.9; Komisarczyk et al., 2024) to calculate exact SHapley Additive exPlanations (SHAP) values. SHAP values, only relatively recently applied in a (soil) ecological context (Cha et al., 2021; Wadoux et al., 2023), quantify the contribution of each predictor variable to the model predictions of the response variable for individual observations, and are expressed in the unit of the response variable (*i.e.,* relative abundance). Positive SHAP values indicate that the predictor variable increases the model-predicted response variable (*i.e.,* relative abundance) relative to the model’s baseline prediction, whereas negative SHAP values correspond to a decrease. We then used the shapviz package (v0.10.3 ;Mayer & Stando, 2023) to create importance barplots and beeswarm plots based on individual SHAP values. SHAP importance barplots show the relative importance of predictor variable, measured as the averaged absolute SHAP value across all observations, thereby reflecting their average contribution to model predictions. SHAP beeswarm plots allow evaluation of the magnitude and direction of each predictor variables’ contribution to the relative abundance for individual observations.

## 3. Results

### 3.1. Sample occurrence and global distribution of fungal nematode antagonists

After filtering the GlobalFungi database for ‘biome’ and ‘type’ that could potentially harbour nematode-antagonistic fungi (see Methods), 27,932 samples were retained (Figure 1A). Most of the selected samples originate from Asia (47.4%), Europe (17.8%), and North America (17.2%) (Figure 1B). It is noted that large global regions including Central and North Africa, the Middle East, Antarctica, western parts of Asia and Russia are under-sampled, while South America is sparsely sampled. The majority of the samples originate from forest biomes (51.2%), followed by grasslands (16.6%) and croplands (13.5%) (Figure 1C).

Within the 27,932 samples that met our biome and type criteria, 59 nematode-antagonistic taxa were detected. In 82.6% of the samples ≥1 antagonist was found, and 34.4% comprised ≥5 antagonists (Figure 2C). For the aggregated biome data, *Clonostachys rosea* (order Hypocreales) (present in 39.4% of the samples), *Metapochonia bulbillosa* (Hypocreales) (37.7%), *Trichoderma viride* (Hypocreales) (27.0%), *Trichoderma asperellum* (Hypocreales) (27.0%), *Metapochonia suchlasporia* (Hypocreales) (25.3%), and *Acremonium persicinum* (Hypocreales) (20.4%) were the six most common nematode antagonists (Figure 2A). Some antagonists were considerably less common, such as *Nematoctonus pachysporus* (Agaricales) and *Brachyphoris* (previously *Dactylella*) *oviparasitica* (Helotiales) that were only found in respectively 0.5% and 0.3% of the samples (Supplementary Table S1). In croplands, relevant plant-parasitic nematode control perspective, the six most common fungal antagonists were *Clonostachys rosea* (present in 67.3% of samples), *Acremonium persicinum* (48.9%), *Chaetomium globosum* (38.4%) (Sordariales), *Metarhizium anisopliae* (Hypocreales) (33.3%), *Trichoderma asperellum* (32.7%), and *Beauveria bassiana* (Hypocreales) (32.4%) (Figure 2B). Subsequently, we determined the mean relative abundances of these six species in all biomes under investigation, and they typically ranged from 0.02% to 0.6% (Supplementary Figure S1).

**Figure 2:**
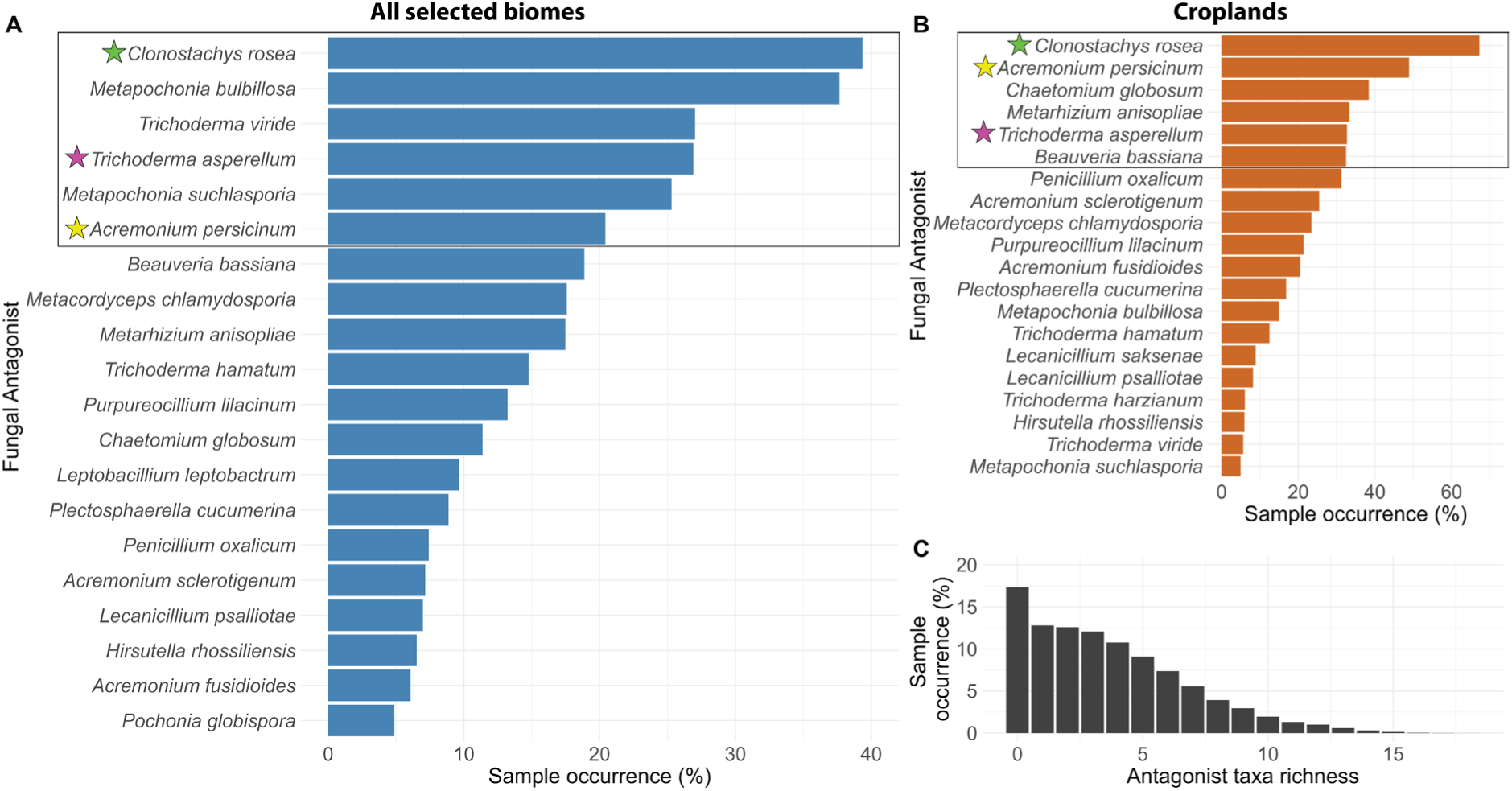
Sample occurrence of fungal nematode antagonists across the GlobalFungi dataset (n=27,932). A) Sample occurrence (in % of samples) for the 20 most common nematode antagonists across all biomes. B) Sample occurrence (in % of samples) for the 20 most common nematode antagonists in croplands (n = 3,774). In both A) and B), black rectangles indicate the six most common fungal antagonists. Coloured stars denote corresponding taxa among the six most common antagonists in all biomes (A) and croplands (B). C) Percentage of samples (Sample occurrence (%)) with an increasing diversity of nematode antagonists.

It is noted that nematode antagonists that are commonly applied as biological control agents, *i.e., M. chlamydosporia*, *P. lilacinum*, *T. harzianum*, *T. viride*, *H. rhossiliensis*, and *Arthrobotrys* spp., were also regularly (5–40% of the samples) detected across all biomes (Figure 2A). While these antagonists did not belong to the highlighted set most common taxa in croplands, they were also regularly detected in this biome (6-23%) (Figure 2B). The mean relative abundance of these antagonists typically ranged from 0.01% to 0.2% in all biomes, although *P. lilacinum* showed higher relative abundances in wetland (14.3%), forest (2.2%), and anthropogenic (1.8%) biomes (Supplementary Figure S2).

The global distribution plots of the six most common fungal antagonists in croplands show that they are widely distributed over the globe and present on all continents (Figure 3). Except for *H. rhossiliensis* (apparently absent from Africa and Antarctica) and *Arthrobotrys* spp. (absent from Antarctica), the six nematode antagonists that are frequently applied as biological control agent also showed a global distribution (Supplementary Figure S3). While *N. pachysporus* and *B. oviparasitica* were uncommon, they showed a wide distribution. *N. pachysporus* was mainly detected in the Americas, Europe and Australia, whereas *B. oviparasitica* was mainly found in the Americas, East Asia and Australia (Supplementary Figure S4). The distribution plots revealed remarkable differences in global distribution between the three *Trichoderma* species. *T. harzianum* was essentially absent from the colder regions of the Northern hemisphere (*i.e*., Canada, Alaska, and Scandinavia) and New Zealand, whereas its sister species *T. viride* and *T. asperellum* showed comparable distributions and were commonly found at these higher latitudes (Figure 3, Supplementary Figures S3 and S5). This suggests that the optimal temperature range of *T. harzianum* is narrower and the preferred temperature higher than those of the sibling species *T. asperellum* and *T. viride*. Acknowledging the substantial differences in sampling intensity between global regions (Figure 1), our analyses show that most fungal nematode antagonists have a widespread distribution at a global scale, and their distribution might even be wider than shown in Figures 3, S3, S4, S5.

**Figure 3:**
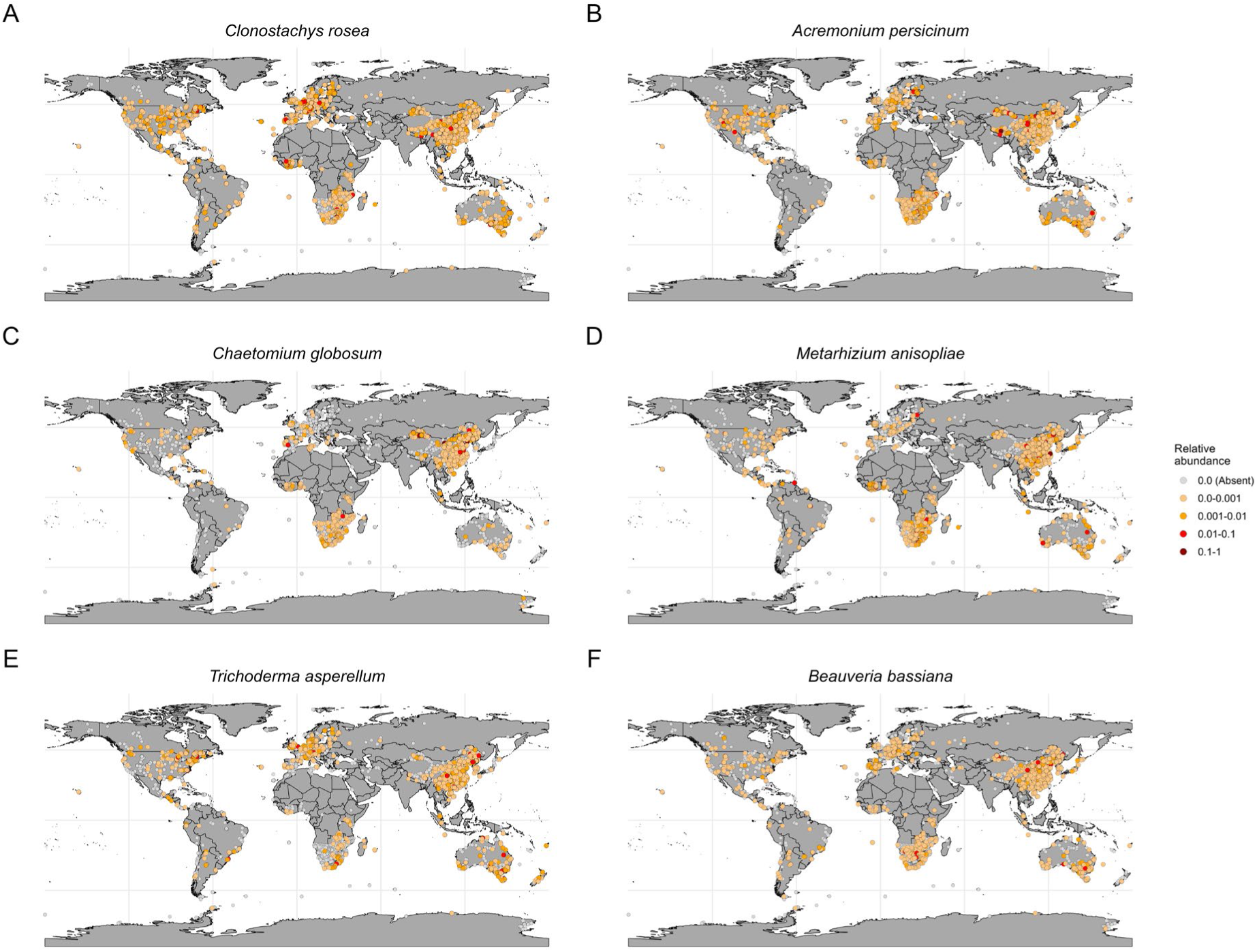
Global distributions of the six most common antagonists in croplands (A: *Clonostachys rosea*; B: *Acremonium persicinum*; C: *Chaetomium globosum*; D: *Metarhizium anisopliae*; E: *Trichoderma asperellum*; F: *Beauveria bassiana*). Coloured dots indicate samples where the taxon was detected; dot color depicts relative abundance. Grey dots indicate samples where the taxon was absent. (n=27,932).

### 3.2. Mode of antagonism frequency

Antagonists having multiple modes-of-action, and antagonists parasitizing eggs & females were most frequently observed, occurring in nearly 70% of the samples (Supplementary Figure S6). In most of those samples more than one taxon with that mode-of-action was present. Furthermore, in 4.9% and 3.0% of the samples, five or more taxa were detected for the modes of antagonism “Multiple, incl. ISR” and “Egg & female parasitic”, respectively. Endoparasitic (25.6%) and nematode-trapping (17.3%) fungi were also common, while taxa relying solely on the production of toxins to predate nematodes were rare (0.5%).

### 3.3. Biome type and climatological conditions affecting nematode antagonist presence and relative abundance

The proportion of samples that contained ≥1 nematode antagonist was highest in anthropogenic (84.6%), cropland (84.5%), and woodland (80.9%) biomes (Figure 4A). A GLMM (binomial) analysis showed that the predicted probability of detecting an antagonist was highest in croplands (88%), whereas woodlands (82%) and grasslands (76%) were not significantly different (Figure 4B). The wide percentile range likely reflects substantial within-biome variation in climatic conditions. As compared to croplands, the relative abundance of the aggregated antagonist community was lower in forests (FST), shrublands (SRL), tundra (TDR) (P<0.001), and wetlands (WTL) (P = 0.002), whereas it was higher in grasslands (GSL) (P<0.001). Using croplands as reference biome, we observed substantial interspecific variation in relative abundances between biomes for the six most common nematode antagonistic taxa in croplands (Figure 5). Some taxa, such as *A. persicinum*, *C. globosum*, and to a lesser extent *M. anisopliae*, showed a higher relative abundance in croplands as compared to most other biomes (Figure 5A, C, D). Other taxa, including the commonly used biological control agents (Supplementary Figure S7), showed more heterogeneous patterns. Notably, *M. chlamydosporia* was generally more abundant in croplands, whereas *T. viride* was consistently more abundant in non-croplands (with deserts and wetlands as an exception).

**Figure 4:**
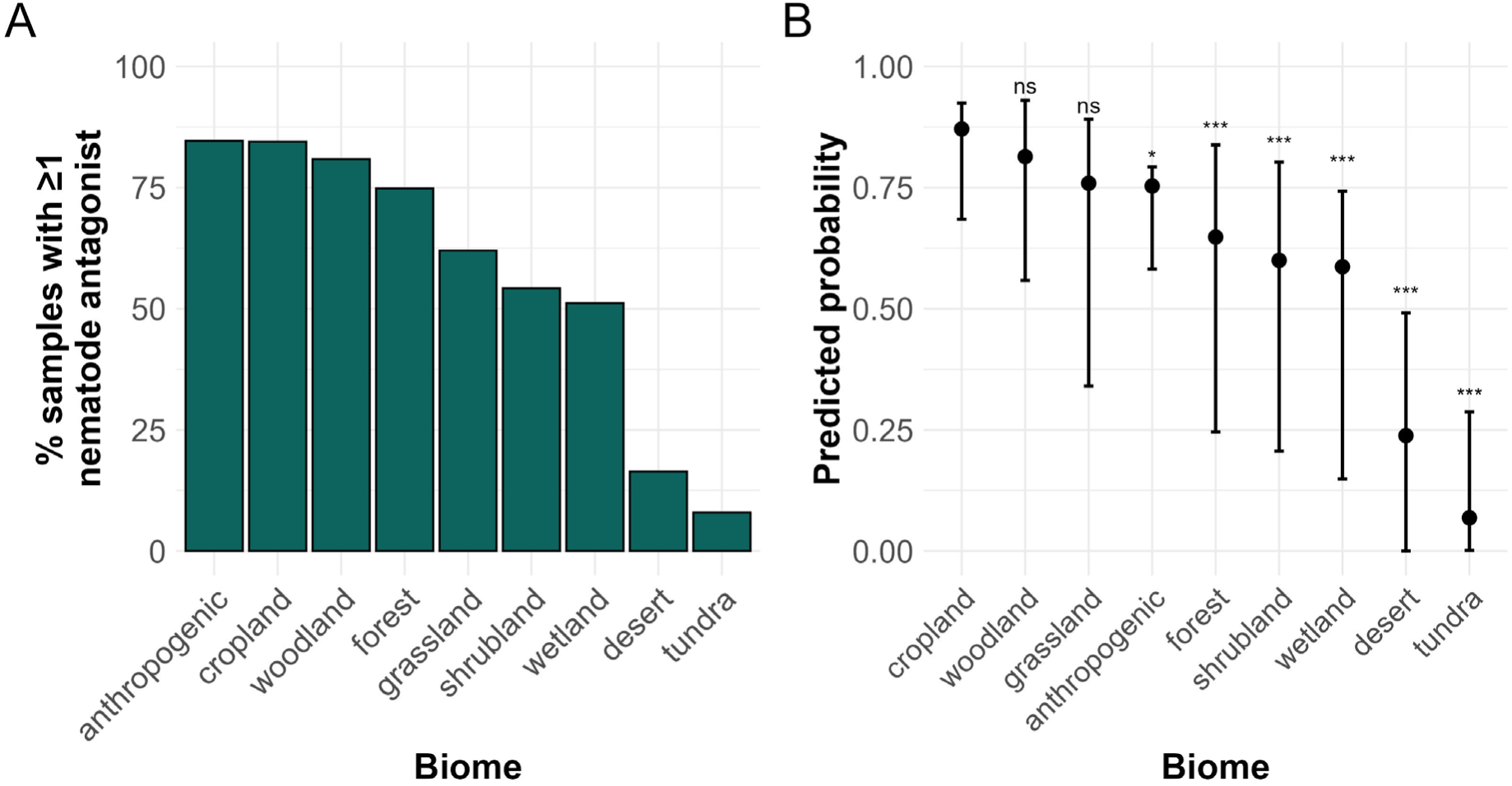
Effect of biome on nematode antagonist presence. A) Percentage of samples in which at least one antagonist was detected, per biome (total n = 27,932). B) Averaged population-level predicted probability of detecting at least one antagonist per biome, based on a GLMM with binomial distribution (total n = 19,450). Asterisks indicate statistical significance as compared to cropland (ns = P > 0.05, * = P < 0.05, and *** = P <0.001). Error bars represent the 2.5^th^-97.5^th^ percentile range of model-predicted probabilities within biomes.

**Figure 5:**
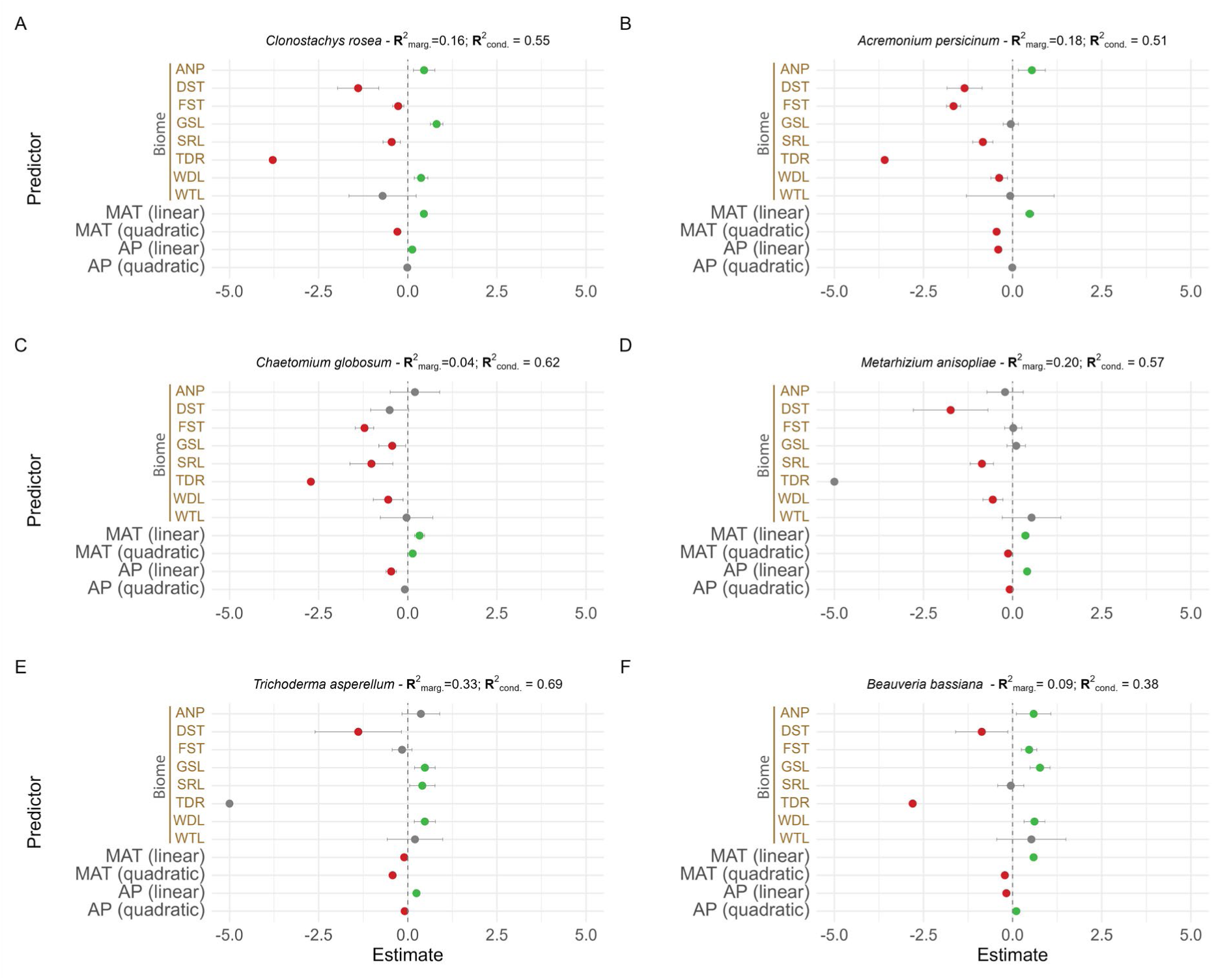
Effect of biomes (as compared to cropland), mean annual temperature (MAT, scaled), and annual precipitation (AP, scaled) on the relative abundance of the six most common fungal nematode antagonists in croplands. Biome abbreviations are: ANP = anthropogenic, DST = desert, FST = forest, GSL = grassland, SRL = shrubland, TDR = tundra, WDL = woodland and WTL = wetland. Model estimates (coloured dots) are shown of GLMMs (total n = 19,450) with a negative binomial distribution, including ‘continent’, ‘month of sampling’, and ‘paper_ID’ as random effects. Dot colour indicates a significant (P*<*0.05) negative (red), significant positive (green), or non-significant (P*>*0.05) (grey) effect. Error bars show 95% confidence intervals. The marginal (*i.e.,* only fixed effects) and conditional (*i.e.,* fixed and random effects) R^2^ of each model is provided at the top of each subplot.

Regarding the impact of temperature and precipitation, both linear (+) and quadratic (-) terms of MAT affected antagonist presence and relative abundance (P<0.001), indicating a unimodal relationship. AP had a significant linear effect on antagonist presence, but no significant effect on the relative abundance of the aggregated antagonist community. Linear and quadratic terms of climatic variables MAT and AP generally affected the relative abundance of the six most common taxa in croplands (Figure 5). *C. rosea*, *A. persicinum*, *M. anisopliae* and *B. bassiana* showed a unimodal response to MAT (Figure 5 A, B, E, F). This implies that their relative abundance increased with MAT until an optimum temperature was exceeded. AP had an antagonist-dependent positive or negative linear effect, whereas its quadratic effect AP was small although occasionally significant (Figure 5 D, E, F). Most of the commonly used biological control agents also showed a non-linear response of their relative abundance to MAT, while their responses to AP differed among taxa (Supplementary Figure S7). A striking difference between the three *Trichoderma* species was observed. Whereas the relative abundance of *T. harzianum* was promoted by higher MAT until an optimum is exceeded, this parameter had a negative effect on *T. viride* and *T. asperellum* (Figure 5E, Supplementary Figure S7C, D). Collectively, these results show heterogenous effects of biome, temperature and precipitation on fungal nematode antagonist presence and relative abundance.

### 3.4. Soil properties affecting relative abundance of fungal antagonists in croplands

We evaluated the importance of individual soil properties (mean ± SD provided in Supplementary Table S2) in explaining the relative abundances of antagonists in the cropland biome. Soil pH contributed most to model predictions of the relative abundance of the aggregated antagonist community (Figure 6A). Low to intermediate pH values overall had negative SHAP values, reflecting lower predicted relative abundances (Figure 6B). In contrast, high pH values, mainly were associated with positive SHAP values, corresponded to increases in predicted relative abundance. For some observations, high pH values increased the model-predicted relative abundance up to 0.09 (≈9% additive effect) relative to baseline prediction. Soil properties CEC, sand content, SOC, and coarse fragments (cfvo) showed a similar importance to one another, whereas clay content had limited predictive value.

**Figure 6:**
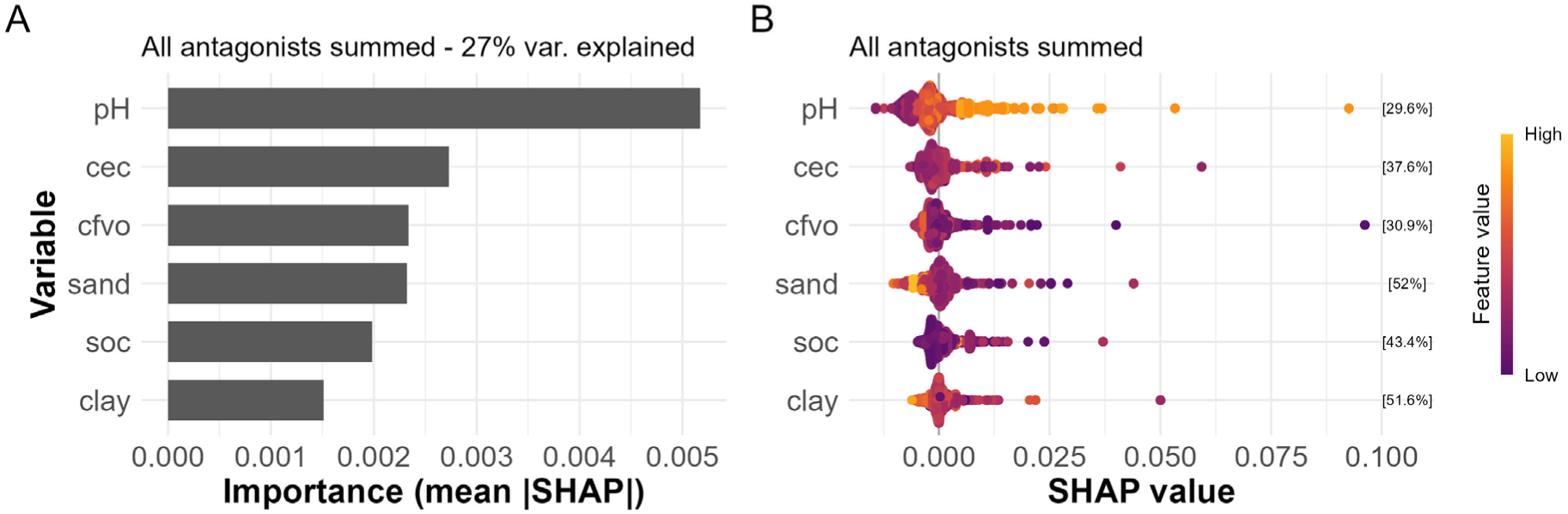
Contribution of soil properties to model-predicted relative abundance of the aggregated antagonist community (*i.e.,* all antagonists summed) across continents (n=2,835). A) Importance barplot showing the mean absolute SHAP value per predictor variable, in decrease order of importance. Higher mean absolute SHAP values indicate higher contributions to model-predicted relative abundance. The percentage above the barplot shows the variation explained by the random forest model. B) Beeswarm plots showing SHAP values of each predictor variable for individual observations (dots), allowing to evaluate directionality of the predictors’ effect. SHAP values larger than 0 correspond to increases in predicted relative abundances, whereas those smaller than 0 correspond to decreases. SHAP values are expressed in the unit of the response (*i.e.*, relative abundance). Colours indicate the relative value of the predictor variable for each individual observation, from low (purple) to high (orange). Percentages between brackets show the percentage of positive SHAP values per predictor variable. cec = cation exchange capacity, cfvo = coarse fragments, sand = sand content (mass %), soc = soil organic carbon, clay = clay content (mass %) (see supplementary Table S2 for details).

Among the six most common antagonists in croplands, the random forest models for *C. rosea*, *A. persicinum*, and *C. globosum* explained >10% of the variation and were therefore used in subsequent SHAP analysis. For *C. rosea,* CEC was the most important variable for model predictions (Figure 7A), with higher CEC values generally being associated with positive SHAP values and lower CEC values with negative SHAP values (Figure 7B). A less pronounced pattern was observed for clay content. Low values of cfvo were generally associated with positive SHAP values. For *A. persicinum*, cfvo was the most important contributors to model predictions (Figure 7C), and had mainly negative SHAP values (87.2%), although some observations showed strong positive contributions (Figure 7D). Higher soil pH values were generally associated with positive SHAP values, whereas lower pH showed the opposite pattern. For *C. globosum,* pH was the most important predictor (Figure 7E), exhibiting a similar pattern as the aggregated antagonist data was observed (Figure 7F). Of the six frequently used nematode antagonists, only *T. harzianum* and *Arthrobotrys* spp. had random forest models explaining more than 10% of the variation (Supplementary Figure S8). For *Arthrobotrys* spp., pH was the most important predictor, also showing a similar pattern as that of the aggregated antagonist community and *C. globosum*.

**Figure 7:**
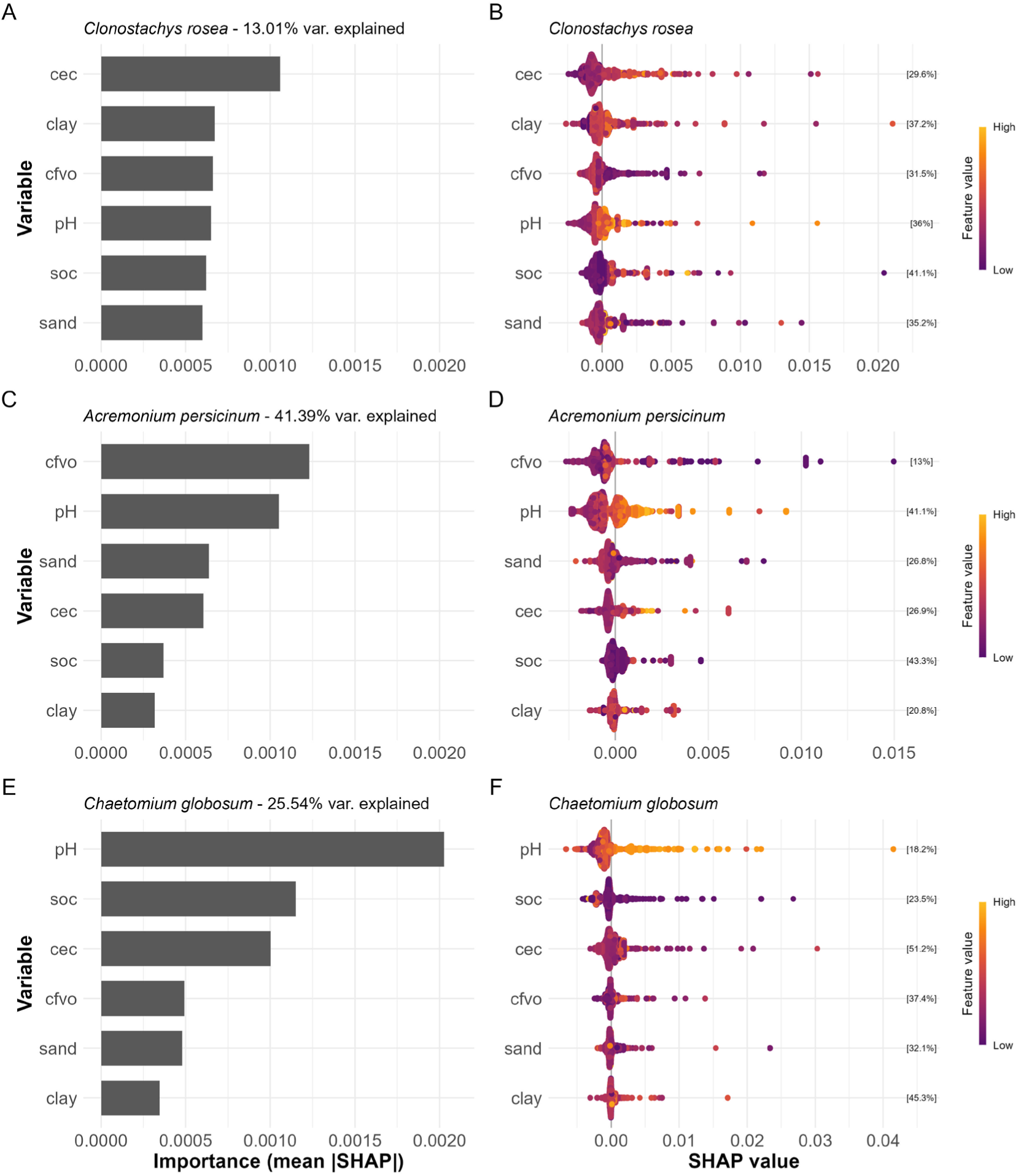
Contribution of soil properties to model-predicted relative abundance of the most prevalent taxa in cropland (across continents) with a model explained variation of more than 10% n=2,835). Panels A, B = *Clonostachys rosea*; Panels C,D = *Acremonium persicinum*; Panels E,F = *Chaetomium globosum*. Left panels: importance barplots showing the mean absolute SHAP values per predictor variable in decrease order of importance. Higher mean absolute SHAP values indicate higher contributions to model-predicted relative abundance. The percentages above the barplots show the variation explained by the random forest model. Right panels: Beeswarm plots showing SHAP values of each predictor variable for individual observations dots), allowing to evaluate directionality of the predictors’ effect. SHAP values larger than 0 correspond to increases in predicted relative abundances, whereas those smaller than 0 correspond to decreases. SHAP values are expressed in the unit of the response (*i.e.*, relative abundance. Colors indicate the relative value of the predictor variable for each individual observation, from low (deep purple) to high (ochre). Percentages between brackets show the percentage of positive SHAP values per predictor variable. Beeswarm plots are scaled ndependently to improve visualization of SHAP distributions. cec = cation exchange capacity, cfvo = coarse fragments, soc = soil organic carbon, sand = sand content (mass %), clay = clay content (mass %).

## 4. Discussion

The global distribution of fungal nematode antagonists, a category of fungi with the potential to play a role in durable management of plant-parasitic nematodes, has not been documented systematically. Using the species-level GlobalFungi database, we retained 28,000 soil samples after filtering for relevant biomes and sample types. We observed a wide distribution of fungal nematode antagonists across the globe. Our analyses revealed that most of the common nematode antagonists show remarkably little biogeography at a global level, and no nematode antagonistic fungus was found to be geographically limited to a single global region. The distribution of individual antagonistic species was affected by biome type and climatic conditions. Nematode antagonists were shown to be remarkably common in croplands, and especially pH was revealed to be an important predictor of relative abundances of nematode antagonists in this biome. Noting that most of these are facultative nematode antagonists that often can live as saprophytes as well, our findings prompt for follow up research on the mechanisms underlying trophic lifestyle switches among nematode antagonists.

### 4.1. Fungal nematode antagonist are abundant and diverse in croplands

We demonstrated that nematode antagonists are remarkably common in both human-impacted and less disturbed soils. Six of the most common nematode antagonists in croplands as well as a selection of six antagonists used as biological control agents were detected on all continents, with possible exceptions of *H. rhossiliensis* (apparently absent from Africa and Antarctica) and *Arthrobotrys* spp. (absent from Antarctica). Antagonist presence was high in croplands and for some antagonist taxa relative abundance was higher in croplands than in relatively undisturbed biomes such as forests and grasslands. This suggests that suitable edaphic conditions for (some) fungal antagonists are present in cropland soils. This observation is consistent with recent studies showing that croplands can support relatively high fungal diversity as compared with undisturbed coniferous or broadleaved woodlands (Labouyrie et al., 2023; Köninger et al., 2023). Possibly contrasting with these observations, croplands are frequently associated with impoverished soil microbial communities (Banerjee et al., 2024; Peng et al., 2024). A study across five European countries revealed that common fungal taxa are enriched in croplands in comparison to adjacent natural grasslands, while the abundance of rare fungi is reduced (Banerjee et al. (2024). Such biotic filtering might explain the presence and abundance of relatively widespread fungal nematode antagonists in croplands.

It is noted that prevalence of nematode antagonists is affected by parameters setting in the analysis. This can be illustrated by *Beauveria bassiana*, a nematophagous and entomopathogenic fungus. Using a minimal threshold of 1,000 reads per sample Větrovský et al., (2019) concluded that the fungus was present in 7.7% of the samples. In our study we only included samples with ≥ 20,000 reads and this thresholds with found *B. bassiana* in 18.9% of the samples. Evidently, a low read number threshold could result in an underestimation of the prevalence of relatively rare fungal taxa. The actual prevalence of antagonists was likely underestimated in our study, as our analysis was restricted to reads that allowed for identification to species level, with an average of 49.3% of the ITS reads meeting this requirement. The detection of fungal taxa may be further influenced by methodological biases inherent to fungal community characterization, such as the choice of primers and DNA extraction efficiency (Lindahl et al., 2013; Reynolds et al., 2022; Schadt and Rosling, 2015).

### 4.2 Common nematode antagonists in croplands

A group of nematode-antagonistic fungi were shown to be common and widespread in croplands, and eight species were detected in more than a quarter of all cropland samples globally. The soil antagonistic community can be steered (Hannula et al., 2021; Cazzaniga et al., 2025; Pasche et al., 2025; Kerry and Crump, 1998) and prevalent antagonistic taxa in croplands may serve as key taxa of interest to induce native soil suppression. However, our understanding of how to effectively steer nematode-antagonistic communities in soils is still in its infancy. It is noted that none of the six most common antagonists in croplands are specialized nematode antagonists. *C. rosea* parasitizes fungi, insects and nematodes (Sun et al., 2020), *A. persicinum* (Liang et al., 2025; Al-Hazmi and Abdul-Razik, 1991) and *C. globosum* (Bairwa et al., 2023; Zhao et al., 2017) are myco- and well as nematode-parasites, *T. asperellum* induces resistance against various plant-pathogenic fungi and nematodes (Fernández et al., 2014; Zheng et al., 2024), and *M. anisopliae* and *B. bassiana* are known non-specific entomopathogens that also feed on nematodes (Karabörklü et al., 2022; Zimmermann, 2007). We conclude that practices associated with arable farming are compatible with the presence of often multiple trophically-generalistic fungal antagonists.

Antagonists from the fungal order Hypocreales were prominently represented in both the aggregated data and cropland-specific subset. In addition to their endophytic and saprophytic lifestyle, many members of the Hypocreales are also characterized as facultative parasites of fungi and invertebrates (mainly insects and nematodes), with parasitic lineages represented in virtually all major hypocrealean families (Kepler et al., 2017). Although some members specifically parasitize fungi (*e.g.*, *Hypomyces odoratus*; Lakkireddy et al., 2020), insects (*e.g., Ophiocordyceps unilateralis*; Kobmoo et al., 2012) or nematodes (*e.g., Hirsutella rhossiliensis*; Chen and Liu, 2005), others appear to parasitize multiple lineages (*e.g., Beauveria bassiana*; McLaughlin, 1962; Karabörklü et al., 2022). The breadth of host preferences among hypocrealean fungi might be explainable by their ability to produce enzymes, such as chitinases, that target conserved structural features shared a across multiple lineages. One such conserved feature is the presence of the polysaccharide chitin, which is found in the fungal cell wall, the nematode eggshells, and insect exoskeletons. As such, the prominence of Hypocreales in soils reflects their ecological versatility in retrieving nutrients from a wide range of sources in the soil environment.

Commercial biological control agents (BCAs) developed to control parasitic nematodes frequently are based on the activity of *P. lilacinum*, *M. chlamydosporia*, *Arthrobotrys,* or antagonistic *Trichoderma* species (Li et al., 2015; Martinez et al., 2023). These taxa were frequently found in both human-impacted and less disturbed soils in our analyses. Despite their potent suppressive capacities in controlled settings (*e.g.*, Hajji et al., 2017; Mukhtar et al., 2021; Loffredo et al., 2024), BCAs are not yet widely used in commercial agriculture. Apart from biological challenges, their adoption is hindered by tedious legislative processes, especially in the European Union (EU). BCAs are regulated similarly to chemical pesticides (*i.e.,* Regulation (EC) No. 1107/2009) and prior to their approval, the EU requests extensive data on the products’ properties, toxicological effects, and impact on biodiversity. Part of these strict regulations for BCAs stems from concerns about introducing non-native organisms in soils. Our data suggests that many of these microbial taxa are naturally widespread in soil, also in the EU, which might justify more lenient EU regulations towards the use of (some) microbial taxa as BCAs in soils.

### 4.3 Climatic conditions affecting fungal nematode antagonists

Global fungal richness and community composition are strongly shaped by climatic conditions (Tedersoo et al., 2014; Větrovský et al., 2019), and we show that climatic factors similarly drive the fungal antagonistic community. We mainly observed non-linear responses of fungal nematode antagonists to climatic conditions, which varied substantially between taxa. Such variation in taxon-specific responses to climatic conditions are frequently observed in studies examining global fungal diversity patterns (Mikryukov et al., 2023; Tedersoo et al., 2014; Van Nuland et al., 2025). The relative abundance of the aggregated antagonist community was not affected by annual precipitation, even though annual precipitation is known to shape fungal soil communities (Tedersoo et al., 2014). Our observation likely reflects taxon-specific responses to annual precipitation among antagonist that could not be captured by a single global model.

Some instances of intra- and intergeneric contrasts in spatial distribution of fungal nematode antagonists have been reported (*e.g.,* Deng et al., 2023a). We observed distinct distributions of three *Trichoderma* species, and *T. harzianum* seemed to have a narrower temperature range and a higher optimal temperature than *T. viride* and *T. asperellum*. An earlier *in vitro* study demonstrated that *T. viride* exhibited stronger antagonistic activity (against *Rhizoctonia solani*) at 10 °C than *T. harzianum*, while this difference was not observed at 25 and 30 °C (Porras et al., 2002). We thus propose that the contrasting latitudinal distributions between *T. viride* and *T. asperellum* on the one hand, and *T. harzianum* on the other hand reflect differences in temperature preference. The potential absence of the endoparasite *H. rhossiliensis* on the African continent is remarkable. *H. rhossiliensis* is the only example of a relatively common nematode antagonist that seems to be absent on a complete continent. The drivers of this apparent geographic exclusion of *H. rhossiliensis* remain unclear. It could be due to the under-representation of north and central Africa soils in the database, or to the obligate nematophagous nature of this fungus (Jaffee et al., 1991).

### 4.4 Soil properties affect nematode antagonists in a species-specific manner

Soil pH was found to be an important predictor of antagonist relative abundance. This finding is corroborated by several studies that have demonstrated the importance of soil pH in shaping soil fungal communities, often in a taxon-specific manner (Liu et al., 2018; Rousk et al., 2010; Li et al., 2025; Tedersoo et al., 2020). For the aggregated antagonist community, *C. globosum*, and *Arthrobotrys* spp., pH values principally at the upper end of the pH gradient were associated with an increase in predicted relative abundance. This pattern suggests a non-linear response of these taxa to pH gradients, where they benefit from an increase in pH once it surpasses a certain threshold. Soil pH of croplands is relatively easy to manipulate through liming (Wenyika et al., 2025). Thus, our analyses suggest altering pH might be a means to steer nematode-antagonistic fungal communities in croplands. It is important to note, however, that these relationships between soil properties and relative abundances are based on model predictions and do not (necessarily) imply causality (Zhang and Wadoux, 2026). Our analysis also revealed that the cation exchange capacity (CEC) of soils was an important predictor variable for the aggregated antagonist community and *C. rosea*. CEC reflects the soil’s potential nutrient retention and its chemical buffering capacity, and is an important driver of soil microbial communities (Docherty et al., 2015; Glassman et al., 2017). A higher CEC generally reflects a more stable soil environment and might favour taxa that are sensitive to rapid changes in soils (Stotzky, 1966), but we don’t expect a narrow relationship between this general parameter and fungal nematode antagonists. In conclusion, these SHAP-based patterns are mainly instrumental for hypothesis formulation (Zhang and Wadoux, 2026) and may give direction to experimental studies testing causality between soil properties and relative abundances of fungal antagonists.

## 5. Conclusion

Our analyses reveal that most fungal nematode antagonists show a widespread global distribution and are present in most soils. Moreover, no fungal nematode antagonist was found which distribution was geographically limited to well-delineated global regions. We demonstrate that (mainly facultative) nematode antagonists are common in soils, this also holds for taxa that are frequently used as main active component in commercial biological control agents. Multiple nematode antagonists were shown to be prevalent and relatively abundant in croplands, implying suitable edaphic conditions for such fungal antagonists in this biome. It should be noted that none of the six most common and widespread antagonists in croplands are exclusive nematode parasites; without exception they also parasitize fungi and/or insects. The prevalence and multiplicity of nematode antagonists in croplands may provide handles to promote plant-parasitic nematode suppressiveness. The perspective of steering of a local nematode antagonist community is attractive as does not require introducing exogenous microorganisms and accompanying legislative challenges. Future studies should focus on pinpointing favourable conditions for (a) an increase in antagonist abundance and (b) for stimulation of a switch towards a nematode-antagonistic lifestyle. (a) Our analyses especially underlined the relevance of pH as a manipulatory soil characteristic to stimulate nematode antagonists in croplands. These results provide leads for future studies aiming to unravel the causal effect of soil properties on nematode antagonist abundance in soils. We furthermore highlight the impact of temperature and - to a lesser extend - precipitation on fungal antagonistic communities. (b) More challenging might be to identify measures that would promote the transition of facultative antagonistic fungi from a mainly saprophytic towards a nematophagous lifestyle under arable farming conditions characterized by relatively nutrient-rich conditions. In this context fundamental insights on the role of arthrobotrisins and related metabolites in trap formation in the model fungus *Arthrobotrys oligospora* (*e.g.,* He et al. 2019) might be instrumental. In summary, our study is the first to systematically determine the global distribution of fungal nematode antagonists. Insights presented here about the commonness and global distribution of a substantial set of nematode antagonists might contribute to exploit the hidden suppressive potential of many soils.

## Author contributions

**R. van Himbeeck:** Conceptualization, Data curation, Formal Analysis, Investigation, Methodology, Software, Visualization, Writing – original draft. **G. Smant:** Conceptualization, Writing – review & editing. **J. Helfenstein:** Conceptualization, Writing – review & editing. **S. Geisen:** Conceptualization, Funding acquisition, Writing – review & editing, Supervision. **J. Helder:** Conceptualization, Funding acquisition, Writing – review & editing, Supervision.

