## Supplementary material for "Global patterns and environmental drivers of soil fungal nematode antagonists"

### 1 Supplementary material

#### *1.1 Supplementary Data S1: R vector containing all queried antagonists.*

c(”Hohenbuehelia auriscalpium”, ”Orbilia auricolor”, ”Orbilia mammillata”, ”Dactylella mammillata”, ”Monacrosporium mammillatum”, ”Acremonium fusidioides”, ”Acremonium persicinum”, ”Acremonium sordidulum”, ”Acremonium strictum”, ”Acremonium sclerotigenum”, ”Acremonium implicatum”, ”Dactylellina haptotyla”, ”Monacrosporium haptotylum”, ”Dactylellina cionopaga”, ”Monacrosporium cionopagum”, ”Monacrosporium ellipsosporum”, ”Orbilia ellipsospora”, ”Dactylella ellipsospora”, ”Dactylellina ellipsospora”, ”Dactylellina parvicolle”, ”Dactylellina phymatopaga”, ”Dactylellina candidum”, ”Dactylella candida”, ”Dactylium candidum”, ”Glomus Intraradices”, ”Glomus mosseae”, ”Glomus versiforme”, ”Glomus fasciculatum”, ”Drechslerella stenobrocha”, ”Harposporium anguillulae”, ”Harposporium cerberi”, ”Harposporium harposporiferum”, ”Podocrella harposporifera”, ”Harposporium arcuatum”, ”Harposporium lilliputanum”, ”Drechmeria coniospora”, ”Drechmeria obovata”, ”Acrostalagmus obovatus”, ”Drechmeria sinensis”, ”Haptocillium sinense”, ”Haptocillium balanoides”, ”Drechmeria balanoides”, ”Verticillium balanoides”, ”Tolypocladium balanoides”, ”Hirsutella rhossiliensis”, ”Hirsutella minnesotensis”, ”Pochonia chlamydosporia”, ”Metacordyceps chlamydosporia”, ”Verticillium chlamydosporia”, ”Cordyceps chlamydosporia”, ”Pochonia globispora”, ”Purpureocillium lilacinum”, ”Paecilomyces lilacinus”, ”Penicillium lilacinum”, ”Lecanicillium psalliotae”, ”Lecanicillium lecanii”, ”Verticillium lecanii”, ”Lecanicillium saksenae”, ”Trichoderma asperellum”, ”Trichoderma citrinoviride”, ”Trichoderma hamatum”, ”Trichoderma harzianum”, ”Trichoderma koningii”, ”Trichoderma tomentosum”, ”Trichoderma virens”, ”Trichoderma viride”, ”Penicillium oxalicum”, ”Chaetomium globosum”, ”Leptobacillium leptobactrum”, ”Hyalorbilia oviparasitica”, ”Brachyphoris oviparasitica”, ”Dactylella oviparasitica”, ”Hyalorbilia fusarina”, ”Vermispora fusarina”, ”Metapochonia suchlasporia”, ”Verticillium suchlasporium”, ”Pochonia suchlasporia", "Metapochonia bulbillosa", "Duddingtonia flagrans", "Plectosphaerella cucumerina", "Nematoctonus concurrens", "Nematoctonus haptocladus", "Nematoctonus leiosporus", "Nematoctonus pachysporus", "Nematoctonus tylosporus", "Nematoctonus subreniformis", "Pleurotus ostreatus", "Coprinus comatus", "Stropharia rugosoannulata", "Catenaria anguillulae", "Catenaria auxiliaris", "Catenaria vermicola", "Haptoglossa heterospora", "Stylopage hadra", "Cystopage cladospora", "Cystopage lateralis", "Rhopalomyces elegans", "Meristacrum asterospermum", "Myzocytiopsis glutinospora", "Myzocytiopsis vermicola", "Myzocytiopsis enticularis", "Myzocytiopsis humicola", "Nematophthora gynophila", "Haptoglossa heterospora", "Haptoglossa zoospora", "Mortierella alpina", "Mortierella globalpina", "Arthrobotrys dendroides", "Arthrobotrys megalospora", "Arthrobotrys iridis", "Arthrobotrys dactyloides", "Arthrobotrys conoides", "Arthrobotrys reticulatus", "Arthrobotrys perpasta", "Arthrobotrys nonseptatus", "Arthrobotrys brochopaga", "Arthrobotrys musiformis", "Arthrobotrys flagrans", "Arthrobotrys ellipsosporus", "Arthrobotrys cylindrosporus", "Arthrobotrys pyriformis", "Arthrobotrys macroides", "Arthrobotrys botryosporus", "Arthrobotrys paucus", "Arthrobotrys polycephalus", "Arthrobotrys sp.", "Arthrobotrys amerosporus", "Arthrobotrys sinensis", "Arthrobotrys koreensis", "Beauveria bassiana", "Metarhizium anisopliae", "Clonostachys rosea")

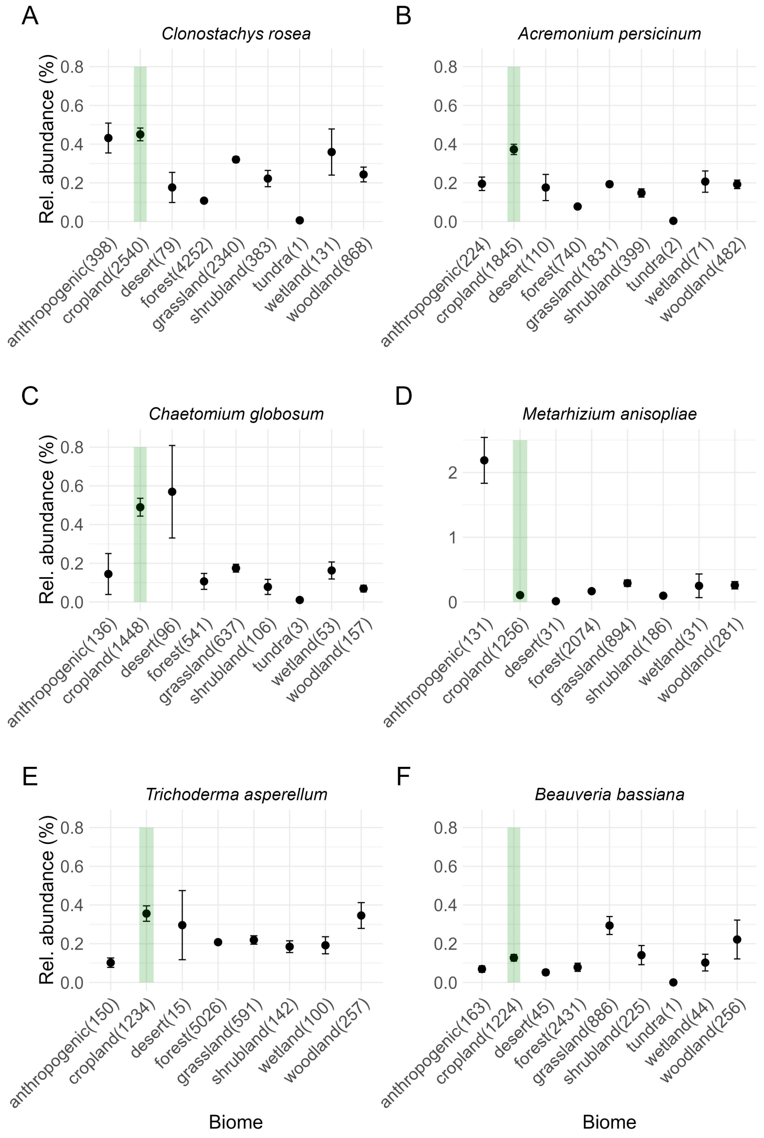


Supplementary Figure S1: Mean relative abundance (black dots, in %) of the six most common taxa in croplands (A: *Clonostachys rosea*; B: *Acremonium persicinum*; C: *Chaetomium globosum*; D: *Metarhizium anisopliae*; E: *Trichoderma asperellum*; F: *Beauveria bassiana*). Only samples where these taxa were detected, were included in the calculation of mean relative abundance. Numbers between parentheses show the number of samples where the taxon was detected in that biome. Error bars show standard deviation. The cropland biome is highlighted by a green rectangle. Biomes where a specific taxon was absent, were omitted from the plots. Note that the y-axis scale for *Metarhizium anisopliae* (D) differs from that of the other taxa.

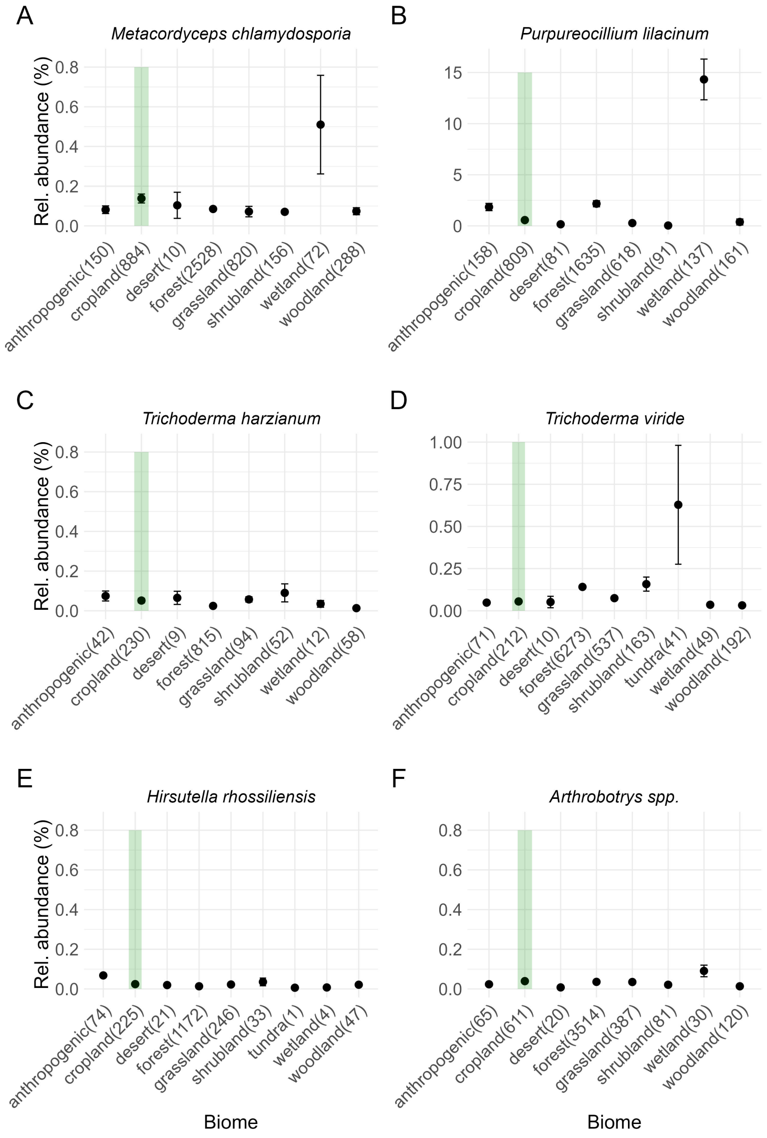


Supplementary Figure S2: Mean relative abundance (black dots, in %) of the six antagonists that are frequently applied as biological control agents (A: *Metacordyceps chlamydosporia*; B: *Purpureocillium lilacinum*; C: *Trichoderma harzianum*; D: *Trichoderma viride*; E: *Hirsutella rhossiliensis*; F: *Arthrobotrys* spp.). Only samples where these taxa were detected, were included in the calculation of mean relative abundance. Numbers between parentheses show the number of samples where the taxon was detected in that biome. The cropland biome is highlighted by a green rectangle. Error bars show standard deviation. Biomes were a specific taxon was absent, were omitted from the plots. Note that the y-axis scales for *P. lilacinum* (B) and *T. viride* (D) differ from those of the other taxa.

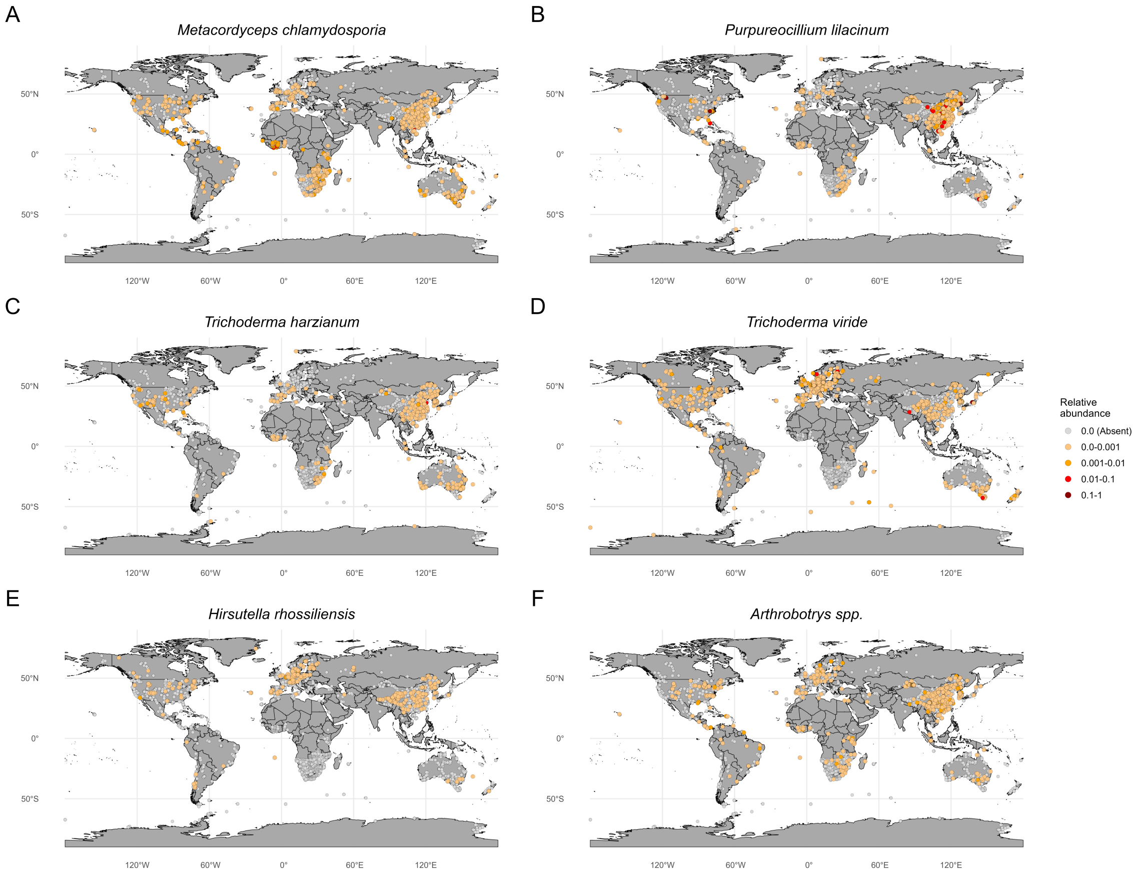


Supplementary Figure S3: Global distributions of the six antagonists that are frequently applied as biological control agents (A: *Metacordyceps chlamydosporia*; B: *Purpureocillium lilacinum*; C: *Trichoderma harzianum*; D: *Trichoderma viride*; E: *Hirsutella rhossiliensis*; F: *Arthrobotrys* spp.). Colored dots indicate samples where the taxon was detected; dot color depicts relative abundance. Grey dots indicate samples where the taxon was absent. (n = 27,932).

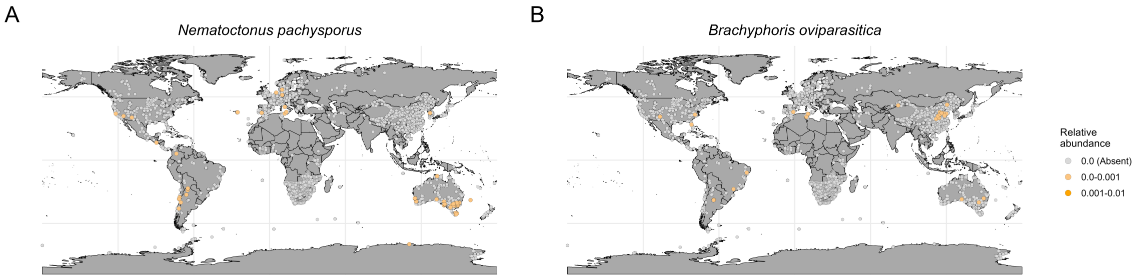


Supplementary Figure S4: Global distributions of *Nematoctonus pachysporus* and *Brachyphoris oviparasitica*. Colored dots indicate samples where the taxon was detected; dot color depicts relative abundance. Grey dots indicate samples where the taxon was absent. (n = 27,932).

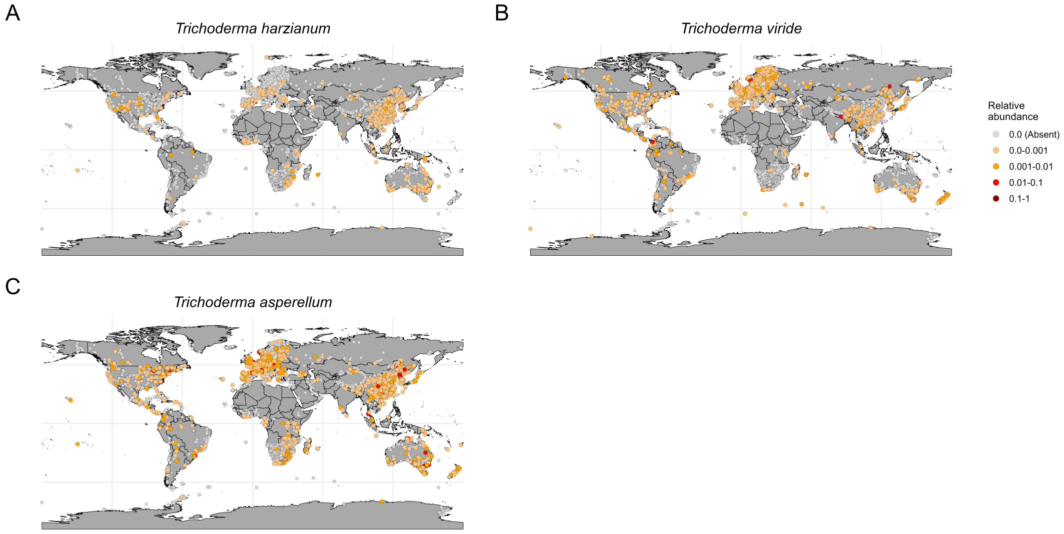


Supplementary Figure S5: Global distributions of *Trichoderma harzianum* (A), *Trichoderma viride* (B), and *Trichoderma asperellum* (C), based on the dataset including samples with >5000 reads (n = 40,926). This distribution plot based on the dataset using less stringent read filtering was included as it more clearly shows the differences in latitudinal distribution between these congeneric fungi. Colored dots indicate samples where the taxon was detected; dot color depicts relative abundance. Grey dots indicate samples where the taxon was absent.

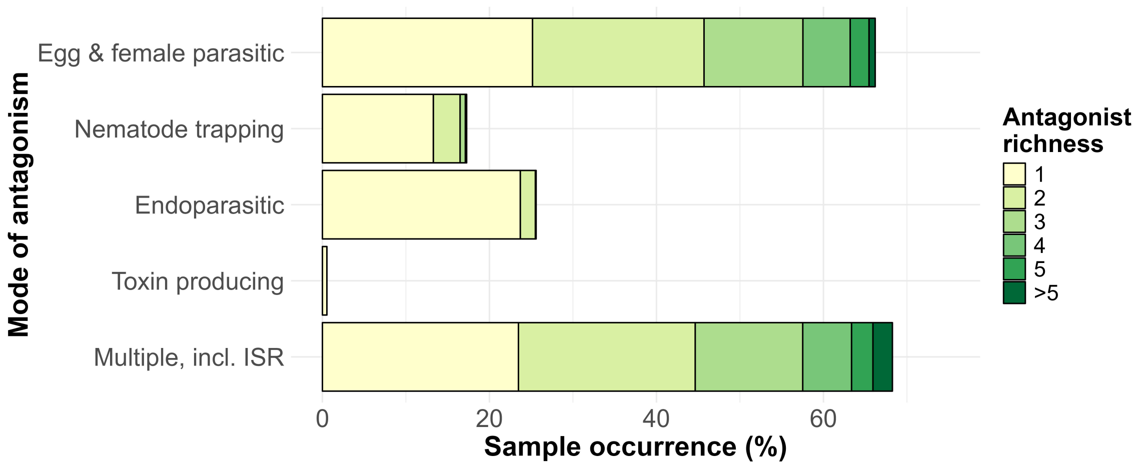


Supplementary Figure S6: Sample occurrence (in % of samples) of each mode of antagonism. Colors indicate the antagonist richness, where occurrences of richness categories larger than 5 were summed. (n = 27,932).

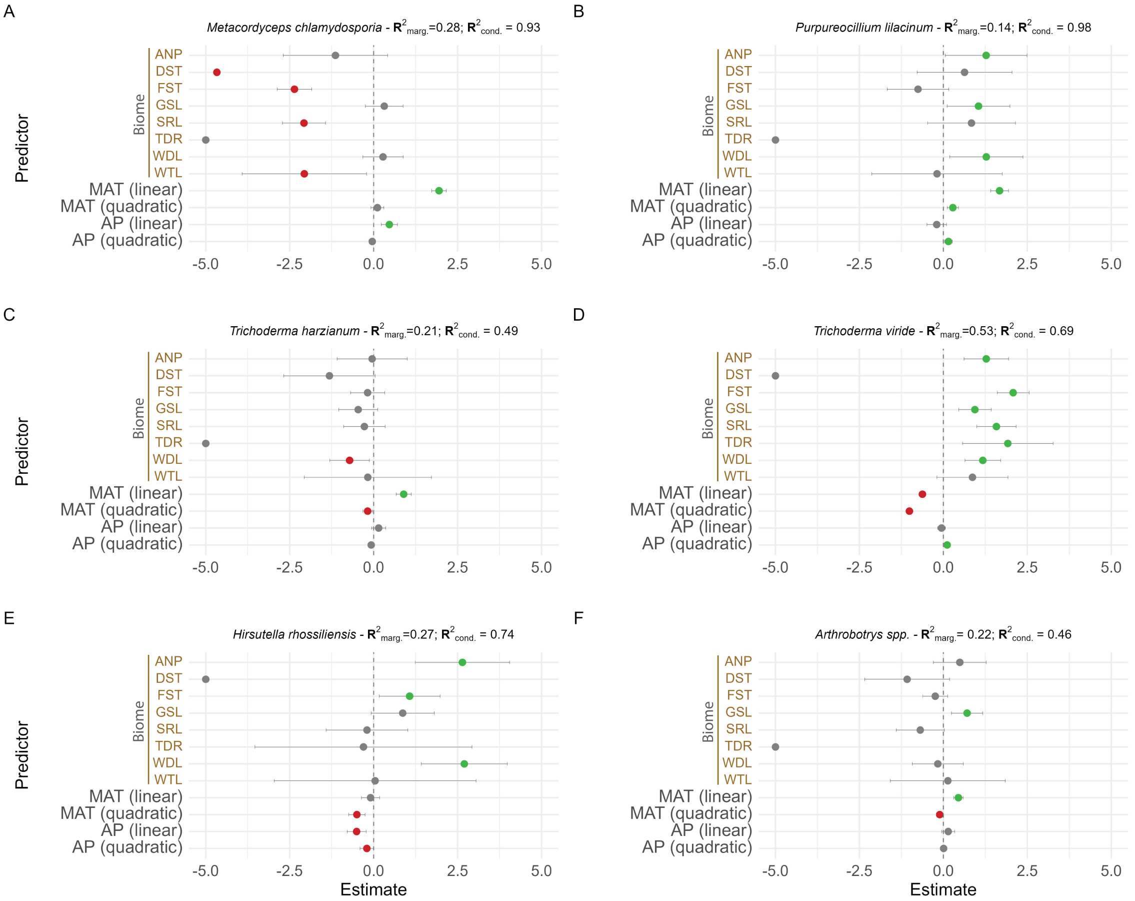


Supplementary Figure S7: Effect of biomes (as compared to cropland), mean annual temperature (MAT, scaled), and annual precipitation (AP, scaled) on the relative abundance of the six antagonists that are frequently applied as biological control agents (A: *Metacordyceps chlamydosporia*; B: *Purpureocillium lilacinum*; C: *Trichoderma harzianum*; D: *Trichoderma viride*; E: *Hirsutella rhossiliensis*; F: *Arthrobotrys spp.*). Biome abbreviations are: ANP = anthropogenic, DST = desert, FST = forest, GSL = grassland, SRL = shrubland, TDR = tundra, WDL = woodland, WTL = wetland. Model estimates (colored dots) are shown of GLMMs (n = 19,450) with a negative binomial distribution, including 'continent', 'month of sampling', and 'paper_id' as random effects. Dot color indicates a significant negative (red), significant ($P<0.05$) positive (green), or non-significant ($P>0.05$) (grey) effect. Error bars show 95% confidence intervals. The marginal (*i.e.,* only fixed effects) and conditional (*i.e.,* fixed and random effects) $R^2$ of each model is provided at the top of each subplot.

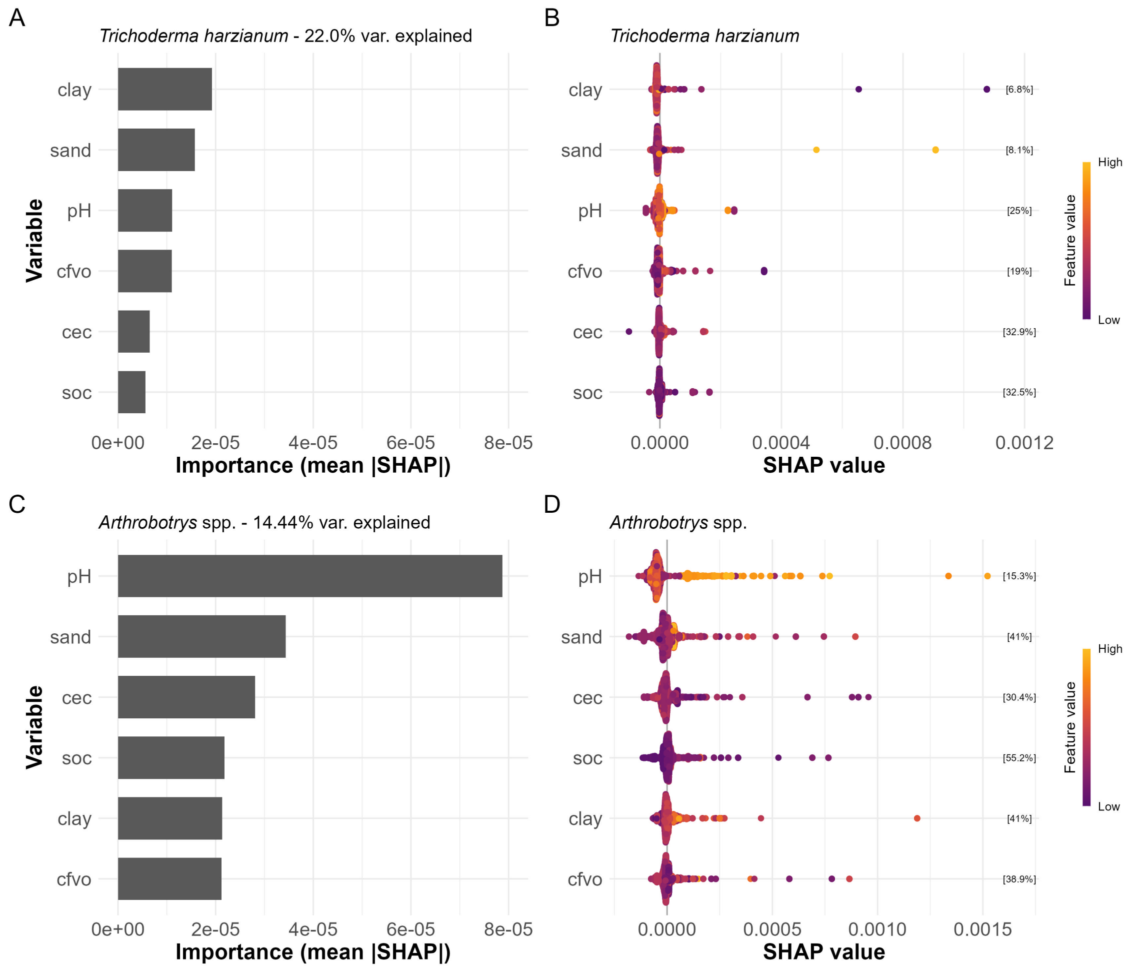


Supplementary Figure S8: Contribution of soil properties to model-predicted relative abundance of the most prevalent taxa in cropland (across continents) with a model explained variation of more than 10% (n=2,835). [A, B = *Trichoderma harzianum*; C,D = *Arthrobotrys* spp.]. A, C) Importance barplot showing the mean absolute SHAP value per predictor variable, in decreasing order of importance. Higher mean absolute SHAP values indicate higher contributions to model-predicted relative abundance. The percentages above the barplots show the variation explained by the random forest model. B,D) Beeswarm plots showing SHAP values of each predictor variable for individual observations (dots). This allows to evaluate the direction (*i.e.,* positive or negative) and effect magnitude for each predictor on model-predicted relative abundances. SHAP values larger than 0 correspond to increases in predicted relative abundances, whereas those smaller than 0 correspond to decreases. SHAP values are expressed in the unit of the response (*i.e.,* relative abundance. Colors indicate the relative value of the predictor variable for each individual observation, from low (purple) to high (orange). Cec = cation exchange capacity, cfvo = coarse fragments, soc = soil organic carbon, sand = sand content (mass %), clay = clay content (mass %).

Supplementary Table S1: Overview of detected antagonists in the filtered GlobalFungi database (n = 27,932), including mode of action, occurrence across all biomes (in % of samples), and exemplary reference of antagonism.

| **Taxon** | **Mode of Antagonism** | **Occurrence (%)** | **Exemplary Reference** |
|---|---|---|---|
| *Clonostachys rosea* | Multiple, incl. ISR | 39.35 | Iqbal et al., 2018 |
| *Metapochonia bulbillosa* | Eggs and/or female parasitic | 37.65 | Moosavi et al., 2010 |
| *Trichoderma viride* | Multiple, incl. ISR | 27.02 | Mukhtar et al., 2021 |
| *Trichoderma asperellum* | Multiple, incl. ISR | 26.90 | Saharan et al., 2023 |
| *Metapochonia suchlasporia* | Eggs and/or female parasitic | 25.30 | Dackman et al., 1989 |
| *Acremonium persicinum* | Multiple, incl. ISR | 20.42 | Al-Hazmi and Abdul-Razik, 1991 |
| *Beauveria bassiana* | Multiple incl. ISR | 18.89 | Fang et al., 2025 |
| *Metacordyceps chlamydosporia* | Eggs and/or female parasitic | 17.57 | Loffredo et al., 2024 |
| *Metarhizium anisopliae* | Endoparasitic | 17.49 | Youssef et al., 2020 |
| *Trichoderma hamatum* | Multiple, incl. ISR | 14.79 | Abd-El-Khair and El-Nagdi, 2014 |

*Table S1 continued*

| **Taxon** | **Mode of Antagonism** | **Occurrence (%)** | **Exemplary Reference** |
|---|---|---|---|
| *Purpureocillium lilacinum* | Eggs and/or female parasitic | 13.21 | Khan and Tanaka, 2023 |
| *Chaetomium globosum* | Eggs and/or female parasitic | 11.37 | Bairwa et al., 2023 |
| *Leptobacillium leptobactrum* | Eggs and/or female parasitic fungi | 9.65 | Regaieg et al., 2011 |
| *Plectosphaerella cucumerina* | Egg and/or female parasitic | 8.87 | Dandurand and Knudsen, 2016 |
| *Penicillium oxalicum* | Multiple, incl. ISR | 7.41 | Martinez-Beringola et al., 2013 |
| *Acremonium sclerotigenum* | Multiple, incl. ISR | 7.15 | Yao et al., 2023 |
| *Lecanicillium psalliotae* | Eggs and/or female parasitic | 6.98 | Krif et al., 2024 |
| *Hirsutella rhossiliensis* | Endoparasitic | 6.53 | Chen and Liu, 2005 |
| *Acremonium fusidioides* | Multiple, incl. ISR | 6.08 | Ogaki et al., 2020 |
| *Pochonia globispora* | Egg and/or female parasitic | 4.88 | Moosavi et al., 2010 |
| *Trichoderma harzianum* | Multiple, incl. ISR | 4.70 | Sahebani and Hadavi, 2008 |

*Table S1 continued*

| **Taxon** | **Mode of Antagonism** | **Occurrence (%)** | **Exemplary Reference** |
|---|---|---|---|
| *Lecanicillium saksenae* | Eggs and/or female parasitic | 4.46 | Sreeja et al., 2025 |
| *Arthrobotrys sp.* | Nematode trapping | 3.30 | Scholler et al., 1999 |
| *Arthrobotrys musiformis* | Nematode trapping | 3.12 | Scholler et al., 1999 |
| *Arthrobotrys megalospora* | Nematode trapping | 2.82 | Scholler et al., 1999 |
| *Mortierella globalpina* | Nematode trapping | 2.76 | DiLegge et al., 2019 |
| *Haptocillium sinense* | Endoparasitic | 2.57 | Zare and Gams, 2001 |
| *Orbilia mammillata* | Nematode traping | 2.52 | Gray, 1984 |
| *Orbilia ellipsospora* | Nematode trapping | 2.14 | Kojima and Saikawa, 2002 |
| *Trichoderma citrinoviride* | Multiple, incl. ISR | 2.02 | Fan et al., 2020 |
| *Arthrobotrys sinensis* | Nematode trapping | 1.37 | Scholler et al., 1999 |
| *Arthrobotrys dactyloides* | Nematode trapping | 0.79 | Scholler et al., 1999 |
| *Arthrobotrys amerosporus* | Nematode trapping | 0.67 | Scholler et al., 1999 |

*Table S1 continued*

| **Taxon** | **Mode of Antagonism** | **Occurrence (%)** | **Exemplary Reference** |
|---|---|---|---|
| *Mortierella alpina* | Multiple, incl. ISR | 0.63 | Al-Shammari et al., 2013 |
| *Pleurotus ostreatus* | Toxin producing | 0.54 | Nyangwire et al., 2024 |
| *Arthrobotrys reticulatus* | Nematode trapping | 0.51 | Scholler et al., 1999 |
| *Nematoctonus pachysporus* | Endoparasitic | 0.49 | Barron and Dierkes, 1977 |
| *Hohenbuehelia auriscalpium* | Nematode trapping | 0.44 | Consiglio et al., 2018 |
| *Acremonium sordidulum* | Multiple, incl. ISR | 0.36 | Schuster and Sikora, 1992 |
| *Drechmeria balanoides* | Endoparasitic | 0.35 | Hay, 1995 |
| *Arthrobotrys flagrans* | Nematode trapping | 0.33 | Scholler et al., 1999 |
| *Arthrobotrys iridis* | Nematode trapping | 0.30 | Scholler et al., 1999 |
| *Arthrobotrys conoides* | Nematode trapping | 0.27 | Scholler et al., 1999 |
| *Brachyphoris oviparasitica* | Egg and/or female parasitic | 0.25 | Becker et al., 2024 |
| *Coprinus comatus* | Nematode trapping | 0.23 | Luo et al., 2007 |

*Table S1 continued*

| **Taxon** | **Mode of Antagonism** | **Occurrence (%)** | **Exemplary Reference** |
|---|---|---|---|
| *Arthrobotrys koreensis* | Nematode trapping | 0.22 | Scholler et al., 1999 |
| *Stropharia rugosoannulata* | Nematode trapping | 0.15 | Zouhar et al., 2013 |
| *Harposporium cerberi* | Endoparasitic | 0.13 | Hodge et al., 1997 |
| *Trichoderma virens* | Multiple, incl. ISR | 0.11 | Herrera-Parra et al., 2017 |
| *Arthrobotrys brochopaga* | Nematode trapping | 0.10 | Scholler et al., 1999 |
| *Orbilia auricolor* | Nematode trapping | 0.06 | Pfister and Liftik, 1995 |
| *Arthrobotrys macroides* | Nematode trapping | 0.05 | Scholler et al., 1999 |
| *Arthrobotrys botryosporus* | Nematode trapping | 0.02 | Scholler et al., 1999 |
| *Catenaria anguillulae* | Endoparasitic | 0.02 | Singh et al., 2012 |
| *Harposporium harposporiferum* | Endoparasitic | 0.01 | Chaverri et al., 2005 |
| *Arthrobotrys polycephalus* | Nematode trapping | 0.01 | Scholler et al., 1999 |
| *Nematoctonus tylosporus* | Endoparasitic | 0.004 | Barron and Dierkes, 1977 |

*Table S1 continued*

| **Taxon** | **Mode of Antagonism** | **Occurrence (%)** | **Exemplary Reference** |
|---|---|---|---|
| *Arthrobotrys dendroides* | Nematode trapping | 0.004 | Scholler et al., 1999 |
| *Arthrobotrys perpasta* | Nematode trapping | 0.004 | Scholler et al., 1999 |

Supplementary Table S2: Mean values and standard deviations of extracted soil properties of the cropland samples (n=2,835).

| **Soil property** | **Mean ± SD** | **Unit** |
|---|---|---|
| SOC | $263.52 \pm 193.91$ | g/kg |
| pH | $6.64 \pm 1.09$ | – |
| cfvo | $1.04 \pm 0.56$ | $cm^3/100cm^3$ |
| CEC | $2.01 \pm 0.55$ | cmol(c)/kg |
| sand | $35.19 \pm 15.84$ | g/100g (%) |
| clay | $26.71 \pm 8.61$ | g/100g (%) |

#### References Table S1


Abd-El-Khair, H. and El-Nagdi, W. M. (2014). Field Application of Bio-Control Agents for Controlling Fungal Root Rot and Root-Knot Nematode in Potato. *Archives of Phytopathology and Plant Protection* 47.10, pp. 1218–1230. ISSN: 0323-5408. DOI: 10.1080/03235408.2013.837632.

Bairwa, A. et al. (2023). Chaetomium Globosum KPC3: An Antagonistic Fungus Against the Potato Cyst Nematode, Globodera Rostochiensis. *Current Microbiology* 80.4, p. 125. ISSN: 1432-0991. DOI: 10.1007/s00284-023-03228-w.

Barron, G. L. and Dierkes, Y. (1977). Nematophagous Fungi: Hohenbuehelia, the Perfect State of Nematoctonus. *Canadian Journal of Botany* 55.24, pp. 3054–3062. ISSN: 0008-4026. DOI: 10.1139/b77-345.

Becker, J. S., Ruegger, P. M., Borneman, J., and Becker, J. O. (2024). Indigenous Populations of a Biological Control Agent in Agricultural Field Soils Predicted Suppression of a Plant Pathogen. *Phytopathology®* 114.2, pp. 334–339. ISSN: 0031-949X. DOI: 10.1094/PHYTO-07-23-0221-R.

Chaverri, P., Samuels, G. J., and Hodge, K. T. (2005). The Genus Podocrella and Its Nematode-Killing Anamorph Harposporium. *Mycologia* 97.2, pp. 433–443. ISSN: 0027-5514. DOI: 10.1080/15572536.2006.11832819.

Chen, S. and Liu, X. (2005). Control of the Soybean Cyst Nematode by the Fungi *Hirsutella Rhossiliensis* and *Hirsutella Minnesotensis* in Greenhouse Studies. *Biological Control* 32.2, pp. 208–219. ISSN: 1049-9644. DOI: 10.1016/j.biocontrol.2004.09.013.

Consiglio, G., Setti, L., and Thorn, R. (2018). New Species of Hohenbuehelia, with Comments on the Hohenbuehelia Atrocoerulea – Nematoctonus Robustus Species Complex. *Persoonia : Molecular Phylogeny and Evolution of Fungi* 41, pp. 202–212. ISSN: 0031-5850. DOI: 10.3767/persoonia.2018.41.10.

Dackman, C., Chet, I., and Nordbring-Hertz, B. (1989). Fungal Parasitism of the Cyst Nematode Heterodera Schachtii: Infection and Enzymatic Activity. *FEMS Microbiology Ecology* 5.3, pp. 201–208. ISSN: 0168-6496. DOI: 10.1111/j.1574-6968.1989.tb03694.x.

Dandurand, L.-M. and Knudsen, G. (2016). Effect of the Trap Crop Solanum Sisymbriifolium and Two Biocontrol Fungi on Reproduction of the Potato Cyst Nematode, Globodera Pallida. *Annals of Applied Biology* 169.2, pp. 180–189. ISSN: 1744-7348. DOI: 10.1111/aab.12295.

DiLegge, M. J., Manter, D. K., and Vivanco, J. M. (2019). A Novel Approach to Determine Generalist Nematophagous Microbes Reveals Mortierella Globalpina as a New Biocontrol Agent against Meloidogyne Spp. Nematodes. *Scientific Reports* 9.1, p. 7521. ISSN: 2045-2322. DOI: 10.1038/s41598-019-44010-y.

Fan, H., Yao, M., Wang, H., Zhao, D., Zhu, X., Wang, Y., Liu, X., Duan, Y., and Chen, L. (2020). Isolation and Effect of Trichoderma Citrinoviride Snef1910 for the Biological Control of Root-Knot Nematode, Meloidogyne Incognita. *BMC Microbiology* 20.1, p. 299. ISSN: 1471-2180. DOI: 10.1186/s12866-020-01984-4.

Fang, M., Wei, X., Sun, J., Wang, A., Tang, H., Wang, L., Leite, L. G., Rasmann, S., Li, J., and Ruan, W. (2025). Management of *Meloidogyne Incognita* with the Endophytic Fungus *Beauveria Bassiana*. *Pest Management Science* 81.9, pp. 5774–5783. ISSN: 1526-498X, 1526-4998. DOI: 10.1002/ps.8948.

Gray, N. F. (1984). The Effect of Fungal Parasitism and Predation on the Population Dynamics of Nematodes in the Activated Sludge Process. *Annals of Applied Biology* 104.1, pp. 143–149. ISSN: 1744-7348. DOI: 10.1111/j.1744-7348.1984.tb05596.x.

Hay, F. S. (1995). Endoparasites Infecting Nematodes in New Zealand. *New Zealand Journal of Botany* 33.3, pp. 401–407. ISSN: 0028-825X. DOI: 10.1080/0028825X.1995.10412966.

Al-Hazmi, A. S. and Abdul-Razik, A. T. (1991). Evaluation of Some Fungal Species as Biocontrol Agents of Meloidogyne Javanica.

Herrera-Parra, E., Cristóbal-Alejo, J., Ramos-Zapata, J. A., Herrera-Parra, E., Cristóbal-Alejo, J., and Ramos-Zapata, J. A. (2017). Trichoderma Strains as Growth Promoters in Capsicum Annuum and as Biocontrol Agents in Meloidogyne Incognita. *Chilean journal of agricultural research* 77.4, pp. 318–324. ISSN: 0718-5839. DOI: 10.4067/S0718-58392017000400318.

Hodge, K. T., Viaene, N. M., and Gams, W. (1997). Two Harposporium Species with Hirsutella Synanamorphs. *Mycological Research* 101.11, pp. 1377–1382. ISSN: 09537562. DOI: 10.1017/S0953756297004152.

Iqbal, M., Dubey, M., McEwan, K., Menzel, U., Franko, M. A., Viketoft, M., Jensen, D. F., and Karlsson, M. (2018). Evaluation of Clonostachys Rosea for Control of Plant-Parasitic Nematodes in Soil and in Roots of Carrot and Wheat. *Phytopathology®* 108.1, pp. 52–59. ISSN: 0031-949X. DOI: 10.1094/PHYTO-03-17-0091-R.

Khan, M. and Tanaka, K. (2023). Purpureocillium Lilacinum for Plant Growth Promotion and Biocontrol against Root-Knot Nematodes Infecting Eggplant. *PLOS ONE* 18.3, e0283550. ISSN: 1932-6203. DOI: 10.1371/journal.pone.0283550.

Kojima, E. and Saikawa, M. (2002). Time-Lapse Photomicrography and Electron Microscopy on Initiation of Infection of Nematodes by Dactylella Ellipsospora. *Mycoscience* 43.4, pp. 299–305.

Krif, G., Lahlali, R., El Aissami, A., Laasli, S.-E., Mimouni, A., Dababat, A. A., Zoubi, B., and Mokrini, F. (2024). Potential Effects of Nematophagous Fungi Against Meloidogyne Javanica Infection of Tomato Plants Under in Vitro and in Vivo Conditions. *Journal of Crop Health* 76.4, pp. 829–839. ISSN: 2948-2658. DOI: 10.1007/s10343-024-00989-7.

Loffredo, A., Edwards, S., Ploeg, A., and Becker, J. O. (2024). Performance of Biological and Chemical Nematicides in Different Soils to Control Meloidogyne Incognita in Tomato Plants. *Journal of Phytopathology* 172.1, e13250. ISSN: 1439-0434. DOI: 10.1111/jph.13250.

Luo, H., Liu, Y., Fang, L., Li, X., Tang, N., and Zhang, K. (2007). Coprinus Comatus Damages Nematode Cuticles Mechanically with Spiny Balls and Produces Potent Toxins To Immobilize Nematodes. *Applied and Environmental Microbiology* 73.12, pp. 3916–3923. DOI: 10.1128/AEM.02770-06.

Martinez-Beringola, M., Salto, T., Vázquez, G., Larena, I., Melgarejo, P., and De Cal, A. (2013). Penicillium Oxalicum Reduces the Number of Cysts and Juveniles of Potato Cyst Nematodes. *Journal of Applied Microbiology* 115.1, pp. 199–206. ISSN: 1364-5072. DOI: 10.1111/jam.12213.

Moosavi, M.-R., Zare, R., Zamanizadeh, H.-R., and Fatemy, S. (2010). Pathogenicity of *Pochonia* Species on Eggs of *Meloidogyne Javanica*. *Journal of Invertebrate Pathology* 104.2, pp. 125–133. ISSN: 0022-2011. DOI: 10.1016/j.jip.2010.03.002.

Mukhtar, T., Tariq-Khan, M., and Aslam, M. N. (2021). Bioefficacy of Trichoderma Species Against Javanese Root-Knot Nematode, Meloidogyne Javanica, in Green Gram. *Gesunde Pflanzen* 73.3, pp. 265–272. ISSN: 1439-0345. DOI: 10.1007/s10343-021-00544-8.

Nyangwire, B., Ocimati, W., Tazuba, A. F., Blomme, G., Alumai, A., and Onyilo, F. (2024). Pleurotus Ostreatus Is a Potential Biological Control Agent of Root-Knot Nematodes in Eggplant (Solanum Melongena). *Frontiers in Agronomy* 6. ISSN: 2673-3218. DOI: 10.3389/fagro.2024.1464111.

Ogaki, M. B. et al. (2020). Cultivable Fungi Present in Deep-Sea Sediments of Antarctica: Taxonomy, Diversity, and Bioprospecting of Bioactive Compounds. *Extremophiles* 24.2, pp. 227–238. ISSN: 1433-4909. DOI: 10.1007/s00792-019-01148-x.

Pfister, D. H. and Liftik, M. E. (1995). Two Arthrobotrys Anamorphs from Orbilia Auricolor. *Mycologia* 87.5, pp. 684–688. ISSN: 0027-5514. DOI: 10.1080/00275514.1995.12026584.

Regaieg, H., Ciancio, A., Raouani, N. H., and Rosso, L. (2011). Detection and Biocontrol Potential of Verticillium Leptobactrum Parasitizing Meloidogyne Spp. *World Journal of Microbiology and Biotechnology* 27.7, pp. 1615–1623. ISSN: 1573-0972. DOI: 10.1007/s11274-010-0615-0.

Saharan, R., Patil, J. A., Yadav, S., Kumar, A., and Goyal, V. (2023). The Nematicidal Potential of Novel Fungus, Trichoderma Asperellum FbMi6 against Meloidogyne Incognita. *Scientific Reports* 13.1, p. 6603. ISSN: 2045-2322. DOI: 10.1038/s41598-023-33669-z.

Sahebani, N. and Hadavi, N. (2008). Biological Control of the Root-Knot Nematode *Meloidogyne Javanica* by *Trichoderma Harzianum*. *Soil Biology and Biochemistry* 40.8, pp. 2016–2020. ISSN: 0038-0717. DOI: 10.1016/j.soilbio.2008.03.011.

Scholler, M., Hagedorn, G., and Rubner, A. (1999). A Reevaluation of Predatory Orbiliaceous Fungi. II. A New Generic Concept.

Schuster, R. P. and Sikora, R. A. (1992). Influence of Different Formulations of Fungal Egg Pathogens in Alginate Granules on Biological Control of Globodera Pallida.

Al-Shammari, T. A., Bahkali, A. H., Elgorban, A. M., El-Kahky, M. T., and Al-Sum, B. A. (2013). The Use of Trichoderma Longibrachiatum and Mortierella Alpina against Root-Knot Nematode, Meloidogyne Javanica on Tomato. *J. Pure Appl. Microbiol* 7, pp. 199–207.

Singh, K. P., Vaish, S. S., Kumar, N., Singh, K. D., and Kumari, M. (2012). *Catenaria Anguillulae* as an Efficient Biological Control Agent of *Anguina Tritici in Vitro*. *Biological Control* 61.3, pp. 185–193. ISSN: 1049-9644. DOI: 10.1016/j.biocontrol.2012.02.005.

Sreeja, P., Reji Rani, O. P., and Chellappan, M. (2025). *Lecanicillium Saksenae*: A Novel Nematophagous Fungus against Root Knot Nematode, *Meloidogyne Incognita* (Kofoid & White, 1919) from Kerala, India. *Journal of Asia-Pacific Entomology* 28.2, p. 102416. ISSN: 1226-8615. DOI: 10.1016/j.aspen.2025.102416.

Yao, Y., Huo, J., Ben, H., Gao, W., Hao, Y., Wang, W., and Xu, J. (2023). Biocontrol Efficacy of Endophytic Fungus, Acremonium Sclerotigenum, against Meloidogyne Incognita under in Vitro and in Vivo Conditions. *Biologia* 78.11, pp. 3305–3313. ISSN: 1336-9563. DOI: 10.1007/s11756-023-01505-4.

Youssef, M. M. A., El-Nagdi, W. M. A., and Lotfy, D. E. M. (2020). Evaluation of the Fungal Activity of Beauveria Bassiana, Metarhizium Anisopliae and Paecilomyces Lilacinus as Biocontrol Agents against Root-Knot Nematode, Meloidogyne Incognita on Cowpea. *Bulletin of the National Research Centre* 44.1, p. 112. ISSN: 2522-8307. DOI: 10.1186/s42269-020-00367-z.

Zare, R. and Gams, W. (2001). A Revision of Verticillium Section Prostrata. VI. The Genus Haptocillium. *Nova Hedwigia* 73.3-4, pp. 271–292. ISSN: 0029-5035. DOI: 10.1127/nova.hedwigia/73/2001/271.

Zouhar, M., Douda, O., Nováková, J., Doudová, E., Mazáková, J., Wenzlová, J., Ryšánek, P., and Renčo, M. (2013). First Report about the Trapping Activity of Stropharia Rugosoannulata Acanthocytes for Northern Root Knot Nematode. *Helminthologia* 50.2, pp. 127–131. DOI: 10.2478/s11687-013-0120-8.
